# Vicariance and dispersal shape the colonization history of the Galápagos carpenter bee, *Xylocopa darwini*

**DOI:** 10.64898/2026.07.19.739469

**Authors:** Graham C. McLaughlin, Verónica Baquero-Méndez, María José Pozo, María de Lourdes Torres, Brian Hollis

## Abstract

Island colonization and subsequent diversification are often shaped by a complex interplay between organism-specific traits including dispersal capacity and ecological requirements, and the configuration of the landscape over geological time. Here, we investigate the evolutionary history and population genomics of the Galápagos carpenter bee, *Xylocopa darwini*, across six islands of the Galápagos archipelago. Despite the bees presumed strong flight ability, we find evidence of long-term isolation among all islands with the strongest divergence occurring between the eastern and western islands, and no evidence of contemporary gene flow between any island pairs. Our analyses indicate that the initial colonization of the archipelago by *X. darwini* occurred during the early Pleistocene, a period when most current islands except for the youngest islands of Isabela and Fernandina, were subaerial, and part of a large contiguous paleo-landmass. Diversification across the Galápagos islands closely tracks the timing of geological separation of the islands, with carpenter bees from the oldest eastern islands splitting first from the western clade over a million years ago. Subsequent diversification of the western clade tracks the sequential fragmentation of the western, central, and southern islands (Isabela, Santiago, Santa Cruz, and Floreana), rather than the geological age of the islands. Despite being on the oldest island, the bee population on Española appears to have been founded more recently (within the last 100,000 years) from San Cristóbal, providing an exception to diversification tracking paleogeography and is the only example of founding via over-water dispersal among the sampled island populations. Measures of contemporary genetic diversity and runs of homozygosity (ROHs) suggest substantially smaller current population sizes on the southern island of Floreana and the eastern islands of Española and San Cristóbal. These results are not consistent with a simple relationship between island area and genetic diversity and instead likely reflect a combination of evolution in isolation following vicariance, rare founder events following over-water dispersal, and possible episodes of secondary contact.

## 1. Introduction

Since the foundational observations of Darwin (1859) and Wallace (1880), islands have provided unparalleled insights into the patterns and processes of biological diversification. Subsequent developments in population genetics, island biogeography, and phylogeography have further established islands as powerful natural laboratories for investigating evolutionary processes (Avise 2000; Emerson 2002; MacArthur and Wilson 1963, 1967; Whittaker et al. 2017; Wright 1931). Volcanic oceanic archipelagos are particularly informative because their extreme isolation from source populations, ecologically complex landscapes, and repeated geographic subdivision promote both endemism and rapid diversification (Barajas-Barbosa et al. 2020; Brée et al. 2026; Cerca et al. 2023; Losos and Ricklefs 2009; Roell et al. 2021). Separation of small populations across environmentally heterogeneous islands amplifies the effects of genetic drift while offering opportunities for strong local adaptation, driving genomic differentiation among populations. Ultimately, however, patterns of genomic divergence within archipelagos reflect a balance between these diversifying forces and gene flow, which acts to homogenize genetic variation. The outcome of this tension depends heavily on the interaction between an organism’s dispersal capacity and the dynamic geological and climatic processes that alter spatial configuration and connectivity through time (Fernández-Palacios et al. 2016; Gillespie 2015; Heaney 2000; Hirschfeld et al. 2023).

The Galápagos archipelago (GA) provides an exceptional system for investigating the interplay among dispersal, isolation, and diversification in a highly geologically and climatologically dynamic landscape. Located approximately 900km west of continental South America, the archipelago comprises 14 major volcanic islands each separated by at least 5 km of ocean (Snell et al. 1996). This extreme isolation, coupled with stark intra- and inter-island environmental heterogeneity (Barajas-Barbosa et al. 2020; Roell et al. 2021) has fueled iconic evolutionary radiations across diverse taxonomic groups (reviewed in Parent et al. 2008). Yet, rapid diversification is far from universal among Galápagos terrestrial fauna. Several highly vagile lineages display little to no genetic population structure across the archipelago (Huyvaert and Parker 2006; Nims et al. 2007; Santiago-Alcaron et al. 2006). In other cases, population structure is governed by dispersal behavior and ecological traits rather than dispersal ability (Chaves et al. 2022; Levin and Parker 2012). Broadly, an absence of genetic differentiation across islands can be attributed to high ongoing inter-island gene flow or a very recent colonization history that has not allowed enough time for the accumulation of genetic differences (Chaves et al 2012; Bollmer et al. 2006).

We are also only beginning to appreciate how the complex geological history of the Galápagos has impacted the partitioning of genetic variation in the archipelago (Heads and Grehan 2021; Harpp and Geist 2018; Orellana-Rovirosa and Richards 2018). Recent studies of the geochemical and geophysical properties of the islands (Geist et al. 2014) and climatological patterns (Karnauskas et al. 2018) suggest that the present-day islands were not formed from a sequential emergence of isolated landmasses but instead have resulted from fusion of emergent volcanoes followed by fragmentation through time (Geist et al. 2014; Karnauskas et al. 2018). Consequently, classic predictions of island colonization — such as the progression rule, where lineage age tracks island age in linearly formed “stepping-stone” island chains (i.e. Hawai’i: Funk and Wagner 1995; Cowie and Holland 2008; Magnacca and Price 2015) — may be confounded by the spatial separation of islands after their initial emergence (Jordan and Snell 2008; Saxton et al. 2023; Poulakakis et al. 2011; Poulakakis et al. 2020).

Further complicating the inference of these evolutionary histories, recurrent periods of island fission and fusion driven by Pleistocene era sea-level fluctuations occurred long after initial geologically driven island separation (Ali and Aitchison 2014; Weigelt et al. 2016). However, the precise impacts of these cyclic geographic connections on the evolutionary histories of endemic taxa remain poorly understood (Myers et al. 2025; Torres-Carvajal et al. 2021; Vlček et al. 2025; Hirschfeld et al. 2023). Phases of secondary contact and gene flow during island fusions can produce highly discordant phylogenetic histories across different genomic markers (Seehausen et al. 2014; Shaw and Gillespie 2016). Furthermore, even when inter-island gene flow has long ceased, the relatively young ages of many Galápagos lineages (Ali & Fritz 2021; Parent et al. 2008) means that most nascent species exhibit relatively shallow absolute evolutionary divergence. Whether driven by ongoing gene flow, widespread incomplete lineage sorting (ILS), or historic introgression and hybridization, early evolutionary genetic studies in the GA relying on one or a few loci often failed to capture the full extent of intraspecific variation. For example, discordance between genetic markers has muddled the phylogenies of the Galapagos *Hogna* wolf spiders, *Calosoma* caterpillar hunter beetles, Galápagos mockingbirds, Darwin’s finches, and marine iguanas (De Busschere et al. 2015; De Busschere et al. 2010; Hendrickx et al. 2015; Arbogast 2006; Hoeck et al. 2010; Nietlisbach et al. 2013; MacLeod et al. 2015; Lamichhaney et al. 2018)

Genomic approaches have recently begun to resolve these complex evolutionary histories (Cerca 2023). Yet, they have so far been applied to a limited subset of Galápagos taxa, primarily charismatic vertebrates such as reptiles (Caccone 2021; Myers et al. 2025; Paradiso et al. 2025) or birds (Lamichhaney et al. 2015, 2018; Vlček et al. 2025; Chavez et al. 2024; Achundia et al. 2025), some plants (Fernández-Mazuecos et al. 2020; Zapata et al. 2024; Gibson et al. 2021), and land snails (Phillips et al. 2019). Only a single study to our knowledge has investigated a GA endemic insect lineage using genome-scale data (Vangestel et al. 2024). This is despite terrestrial arthropods comprising roughly 97% of the archipelago’s terrestrial animal species diversity (Galapagos Species Database; Peck 2001; Peck 1991). This taxonomic bias skews our broader understanding of Galápagos biogeography as differences in histories, traits related to dispersal, and ecological characteristics across clades inevitably shape colonization timelines, inter-island dispersal dynamics, historical demographics, and localized extinction risks (Avise 2016).

The Galápagos hosts a single endemic bee species (Galápagos Species Database; Ontenada-Gallagos et al. 2026), the large carpenter bee, *Xylocopa darwini* (Cockerell, 1926). This imbalance when compared to the diversity of bees in the neotropics (Michener 2007; Moure et al. 2007) underscores the difficulty of transoceanic dispersal (Gillespie and Roderick 2002). By boring nests deep into dead stems of woody plants, the ancestors of *X. darwini* were uniquely suited for long-distance transoceanic dispersal via rafting (Cockerell 1935; Hanna 1932; Poulsen and Rasmussen 2020; Ali and Fritz 2021). This hypothesis is corroborated by the co-occurrence of a nest-parasitizing beetle obligately associated with *Xylocopa* (Linsley 1966). Recent broad-scale phylogenetic revisions of Neotropical *Xylocopa* indicate that the closest extant mainland relatives – currently distributed from Peru to Brazil – split with the GA lineage between 1 and 4 Mya (Blaimer et al. 2018; Melo and Martins 2025). However, this deep divergence event does not date the true arrival time to the archipelago, particularly if key mainland source populations are extinct or unsampled. A more comprehensive history of colonization requires dating a split derived from the earliest divergence events within the archipelago itself (Ali and Fritz 2024). Currently, both the precise timing and subsequent colonization pathway of *X. darwini* throughout the archipelago remains unresolved, obscuring how historical island configuration and paleogeography have interacted with migration to shape the evolution of *X. darwini* in the GA. Although *X. darwini* is distributed widely across both older islands (*e.g.* San Cristóbal, Española) and younger islands (*e.g.* Isabela, Fernandina, Genovesa), early taxonomic work reported no consistent morphological differentiation among island populations (Cockerell 1935). This contrasts with patterns observed in many other GA taxa and led to the historical hypothesis that ongoing inter-island migration or recent dispersal across the GA has limited evolutionary divergence (Cockerell 1935; Linsley & Usinger 1966). Alternatively, the bee may represent an example of cryptic genetic differentiation, which has been found in other GA insect lineages (Schmitz et al. 2007). This is partially supported by analysis of *X. darwini mt*DNA, which has identified at least two major historically isolated lineages (an eastern and western clade) with non-overlapping haplotype distributions (Vargas et al. 2015). However, extensive haplotype sharing is observed within each of these clades and paraphyletic relationships between individuals from the same island suggest recurrent inter-island migration, recent divergence within island groups, or simply the limited resolution of single-locus data.

To address these gaps and shed light on the diversification of *X. darwini* across the archipelago, we generated the first *de novo* draft reference genome for the Galápagos carpenter bee and conducted whole-genome resequencing of individuals from six of the eleven major islands where extant populations persist (Fig. 1). Additionally, we sampled individuals from four distinct shield volcanoes comprising the island of Isabela (Harpp and Geist 2018). These volcanoes function as isolated, de facto islands for many endemic Galápagos taxa due to the formidable dispersal barriers imposed by surrounding historic lava flows (Arteaga et al. 2019; Pozo et al. 2025; Gentile et al. 2009; Gaughran et al. 2024). Specifically, we aimed to quantify genome-wide differentiation among island populations to evaluate the extent of contemporary gene flow and reconstruct a temporally resolved phylogeographic and demographic hypothesis for *X. darwini*. We then revisit the question of morphological differentiation between island populations by measuring body mass from individuals collected from different sites across multiple sampling seasons. As the sole native bee species in the Galápagos, *X. darwini* likely acts as a critical keystone pollinator that visits a significant proportion of the archipelago’s endemic flora (Chamorro et al. 2012; Linsley 1966; McMullen 1999). Consequently, core population genetic parameters not only clarify ancient colonization dynamics but also establish a baseline for population health and support long-term conservation of *X. darwini* in the GA.

**Figure 1.**
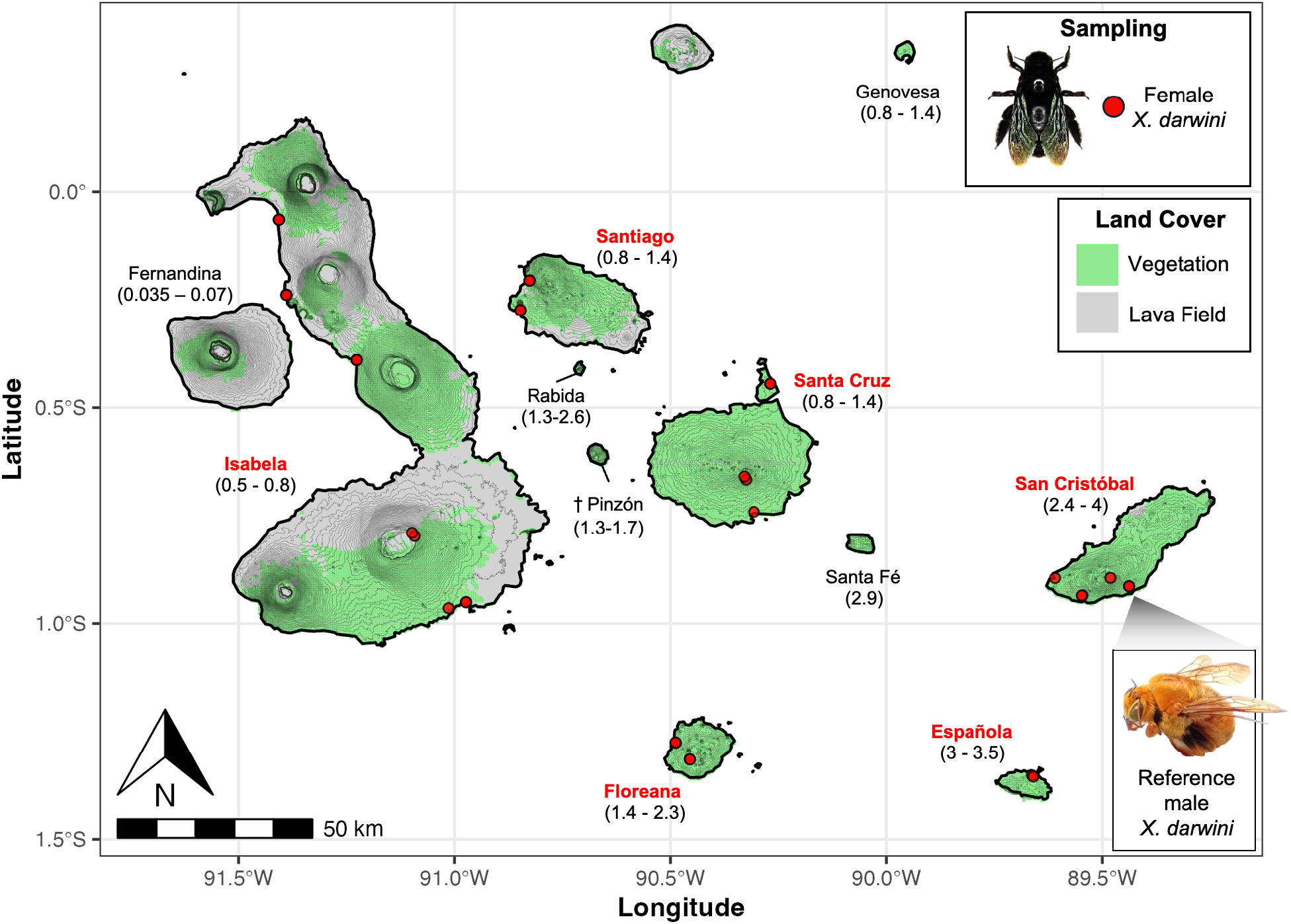
Sampling breadth of the sequenced *X. darwini* individuals. Red circles represent sampling locations of 42 resequenced females and the sampling location of the male used to build the reference genome is also noted. Sampled islands are labeled in red. Islands labeled in black were not sampled in this study but host a known extant population, except for the island of Pinzón for which only museum specimens have been reported (Ontenada-Gallagos et al. 2026). Unlabeled islands host no reported historical or contemporary *X. darwini* population. Island ages from Geist et al. (2014) are reported under each label in millions of years. Island shapefiles come from NOAA GSHHGD (Wessel and Walter, 2025), land cover data comes from NASA MODIS (Friedl and Sulla-Menashe, 2019), and topographic contours come from NASA STRM (Jarvis et al. 2013). Photo of *X. darwini* female taken by Graham C. McLaughlin. Photo of *X. darwini* male taken by Veronica Baquero-Méndez.

## 2. Materials and Methods

### 3. Results

#### 3.1 Reference Genome

We generated 11.36 Gb of PacBio HiFi sequence data comprising 1.08M reads with a mean read accuracy exceeding Q30 (>99%). Mean and median read lengths were 10,640 bp and 9,321 bp, respectively, with read lengths ranging from 105 to 72,979 bp (fig. S1). Mean and median per-base sequencing depth were 53.4× and 55×, respectively (fig. S2). Genome size, estimated from sequencing depth by dividing the total bases sequenced by the mean coverage, was ∼213.1 Mb. In comparison, a k-mer-based estimated using GenomeScope (k = 21) produced a genome size of 184.6Mb.

Among the four assembly algorithms tested (see Materials and Methods) the HiFlye draft assembly had the greatest contiguity and highest BUSCO completeness. This assembly comprised 433 contigs with a total length of 215,970,125 bp (fig. S3). The largest contig was 20,681,599 bp, with an N50 of 11.7 Mbp and an N90 of 276 Kbp (fig. S3). The L50 and L90 were 8 and 26 contigs, respectively (fig. S3). BUSCO analysis recovered 99.7% of expected hymenopteran orthologs, indicating a highly complete assembly (fig. S3).

#### 3.2 Resequencing, Mapping, Variant Calling, and Datasets

Whole-genome resequencing yielded an average of 58M reads per individual (29M reads per direction per individual). The number of total reads per sample ranged from 42M to 88M. After adapter trimming and quality filtering, an average of 98.7% of reads were retained with >99% of bases exceeding a Phred score of Q38 across all samples. After mapping to the reference, the mean per-sample mapping rate was 73.5%, ranging from 62.5% to 84.5% (fig. S4). We tested for reference bias by comparing mapping rate between individuals from separate islands and found no significant decrease in mapping rate among the non-reference western. We graded contigs based on mean mapping quality (MQ), breadth of coverage, and median sequencing depth. We removed 109 contigs with mean MQ < 30, 288 contigs with >90% breadth of coverage, and 12 contigs exhibiting high read depth each. In total, this filtering removed 30,537,182 bp spanning 409 contigs with a mean length of 74,663 bp and a range of 507bp-1.57 Mbp, resulting in a final reference dataset of 185,432,943 sites spanning 24 core contigs used for downstream variant calling. This more liberal contig filtering strategy retains well-supported nuclear regions while excluding low-quality, highly repetitive, or potentially contaminated contigs, such as those derived from endosymbionts.

Before variant filtering, the mean and median per-sample read depth of the GA samples across this reference set was 21.7× and 22.4× respectively (min = 13.2×, max = 29.4×) (fig. S5). The mean per-site missingness rate was ∼0.01 across all GA samples and the per-sample fraction of missing sites ranged from 0.00231 to 0.017 in with a mean of ∼0.01 (fig. S6). The outgroup *X. virginica* sample had a mean read depth of 32.5× and a fraction missingness of ∼0.07. After filtering, we retained a total of 145,507,563 sites containing 3,273,028 SNPs across all Galápagos samples.

#### 3.3 Population Structure

Principal component analysis (PCA) separated the samples into 5 distinct clusters across PC1 and PC2, which together explained 33.51% of the variation present in the 125,664 unlinked SNP dataset (Fig. 2A). PC1 explained the majority of this variation (19.9%) and was defined by large separation in PC space of the westernmost sampled island of Isabela from the easternmost islands of San Cristóbal + Española (*Δ*_PC1_ = 0.404, *p* < 0.0001) with no significant separation of San Cristóbal and Española (*Δ*_PC1_ = 0.0009, *p* = 0.887). The central island cluster of Floreana, Santa Cruz, and Santiago were placed in between Isabela and the eastern islands. Of the central islands, Floreana showed the most similarity to the San Cristóbal-Española cluster (*Δ*_PC1_ = – 0.277, *p* < 0.0001), though there was still significant separation. All central islands showed relatively shallow separation on PC1 (*Δ*_PC1_: 0.005 to 0.088). PC2 again showed no separation between the eastern islands of San Cristóbal and Española (*Δ*_PC2_ = 0.001, *p* = 0.850) with the largest separation being between Santa Cruz and San Cristóbal (*Δ*_PC2_ = 0.469, *p* < 0.0001) and the shallowest significant separation between Santiago and Floreana (*Δ*_PC2_ = 0.014, *p* < 0.0001). A separate PCA including only the four volcano populations on Isabela found no evidence of significant clustering on PC1 or PC2 (Fig. 2A). However, a PCA run only including San Cristóbal and Española did find significant separation of the two eastern islands (*Δ*_PC1_ = 0.8366, *p* < 0.0001) across PC1 of this subsetted analysis (Fig. 2A). Percentages of variance explained by lower PCs are shown in fig. S7.

**Figure 2.**
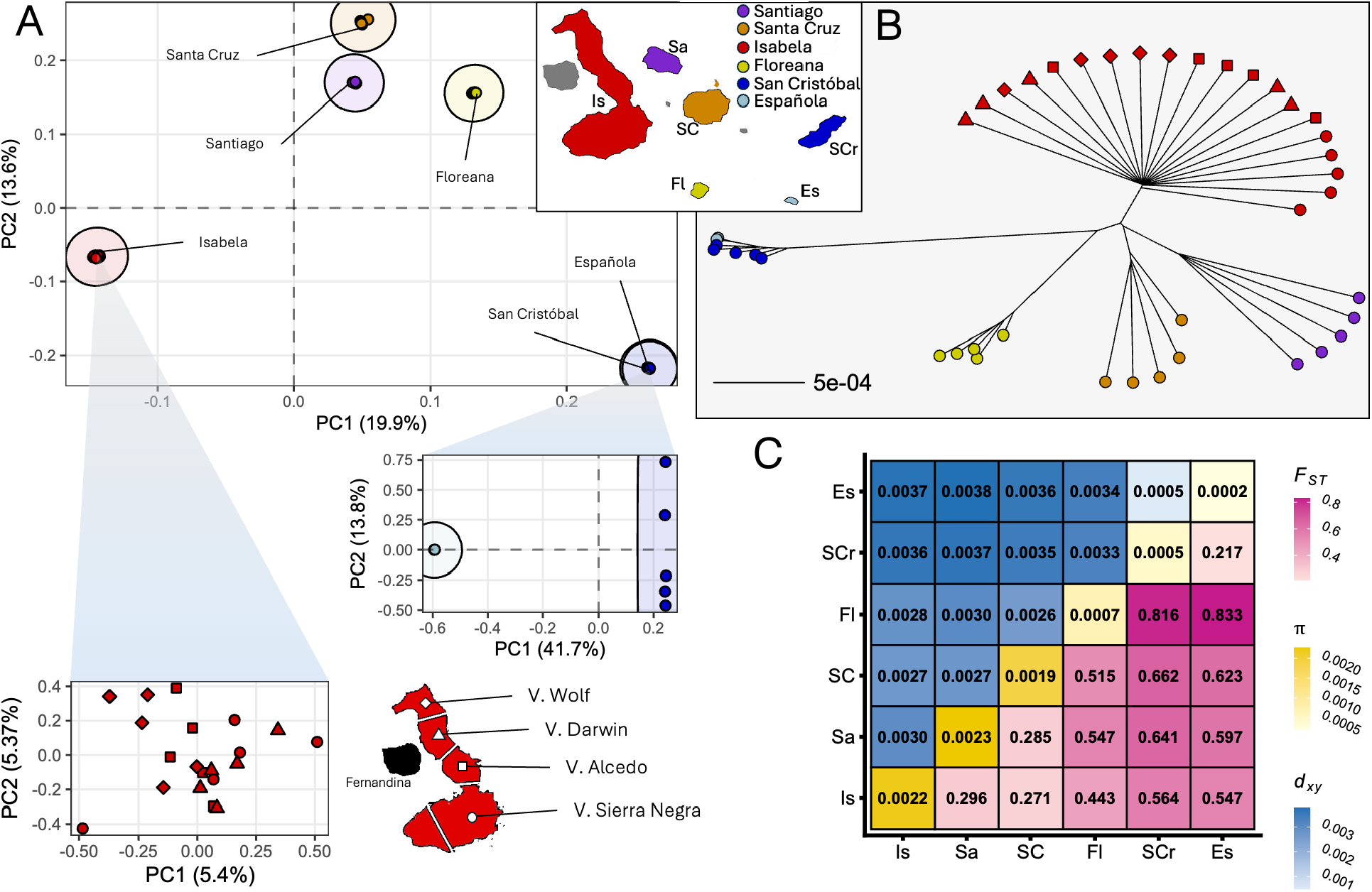
Population structure of the six sampled *X. darwini* island populations. (A) PCA including all 42 resequenced individuals. A separate PCA including only the 20 Isabela individuals is depicted with shapes denoting the volcano-of-origin of each individual. A PCA including only the individuals from Española and San Cristóbal is also shown. (B) Neighbor-joining *d_xy_* tree of all individuals. Shapes again denote the volcano-of-origin of each Isabela individual. (C) Pairwise genetic distance matrix including values of *d_xy_*, within-island nucleotide diversity (π), and relative genetic differentiation (F_ST_).

The individual-level *d_xy_* tree again recovered individuals sampled from the same island as strictly monophyletic and separated by long internal branches (Fig. 2B). Individuals sampled from San Cristóbal and Española formed a reciprocally monophyletic group with individuals from each island tightly clustering together on the tree. Concordant with the PCA, these two clades were separated by a very short branch length. Individuals sampled from the four Isabela volcanoes were paraphyletic with respect to one another, with no consistent grouping among individuals sampled from the same volcano. A long internal branch separated the eastern islands from all others.

Pairwise average F_ST_ differed significantly from zero between all pairs of islands (Fig. 2C) except for the Isabela volcano populations (fig. S8). Significant F_ST_ values ranged from 0.217 between Española-San Cristóbal to 0.816-0.833 between Floreana-San Cristóbal and Floreana-Española. Values between the central islands ranged between 0.271 for Santa Cruz-Isabela to 0.285 for Santa Cruz-Santiago. Values between the central islands and the eastern islands, including Floreana, ranged between 0.443 to 0.662. Within-population nucleotide diversity (π) also differed significantly among island populations, with the southern island of Floreana and especially the eastern islands of Española and San Cristóbal showing greatly diminished nucleotide diversity compared to the central and western islands (Fig. 2C, diagonal).

#### 3.5 Island Relationships and Divergence Times

The species tree inferred from 20 kb-windowed gene trees using ASTRAL-IV recovered two major clades, an eastern island clade consisting of San Cristóbal and Española, and a western/central island clade consisting of Floreana, Isabela, Santa Cruz, and Santiago (fig. S9). Within the western/central island clade, Santa Cruz and Santiago form a monophyletic group that is successively sister to Isabela, and then Floreana. Although all nodes were strongly supported (local posterior probability (PP) = 1), quartet frequencies revealed substantial gene tree discordance in nodes separating the central/western islands, especially between Isabela, Santa Cruz, and Santiago (table S4).

Divergence times estimated using StarBEAST3 with a fixed clock rate of 8.4 x 10^-4^ substitutions site^-1^ Myr^-1^ recovered all island relationships with strong support (PP = 1) (Fig. 3A). We estimated the deepest split among the sampled islands to have occurred between 1.237 and 1.645 Mya (95% Highest posterior density (HPD); mean = 1.429 Mya). Within the western/central clade, the deepest split separates Floreana between 604 and 803 Kya (95% HPD; mean = 698 Kya), followed by Isabela between 399 and 547 Kya (95% HPD; mean = 470 Kya), and finally Santiago and Santa Cruz which diverged between 279 and 412 Kya (95% HPD; mean = 344 Kya). The shallowest split of the sampled islands is between San Cristóbal and Española between 45 and 78 Kya (95% HPD; mean = 61 Kya) (Fig. 3A). Overall, we observe a deep, ancient split separating the eastern and western islands followed by relatively rapid divergence between the western island populations.

**Figure 3.**
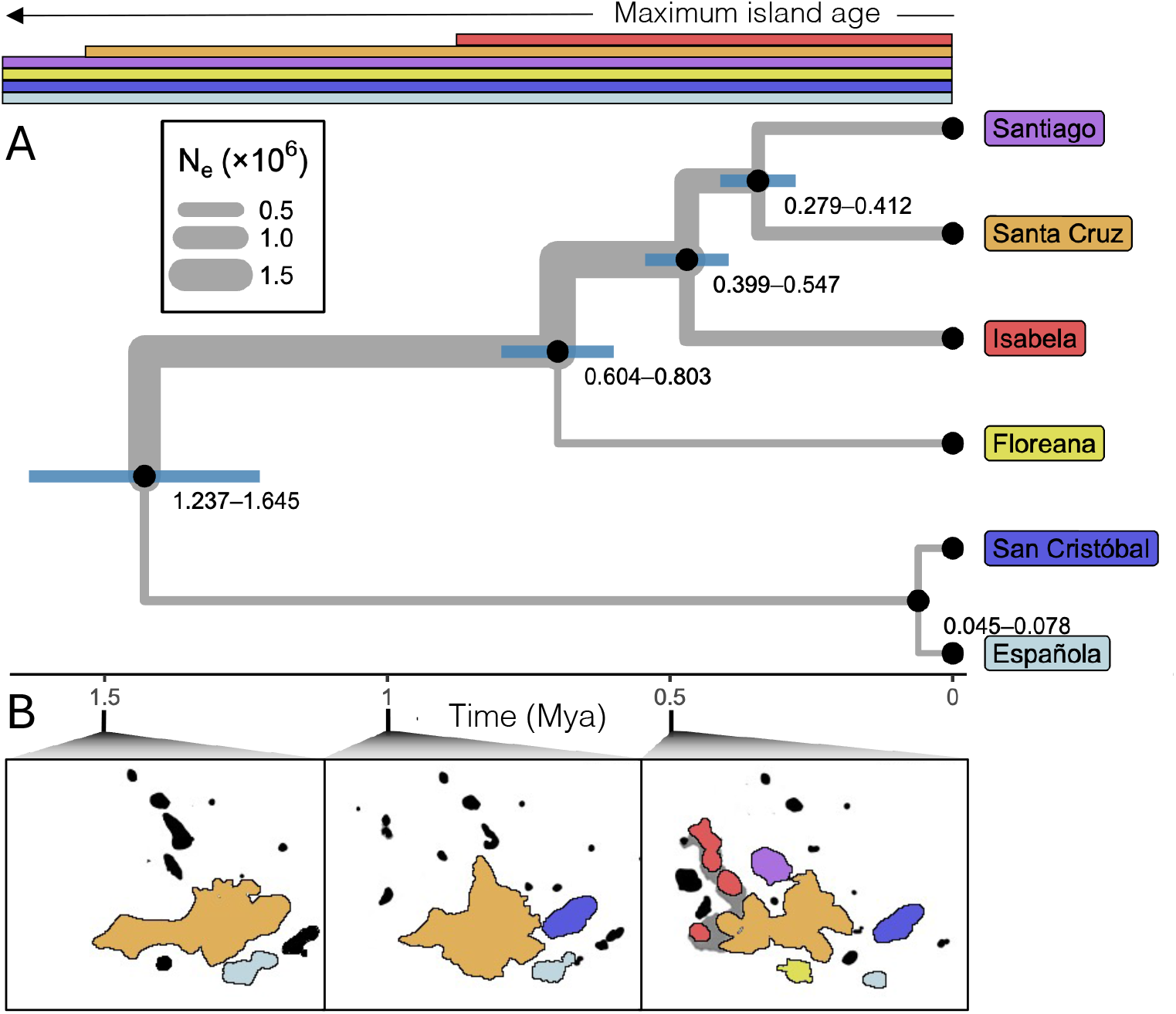
(A) Time-calibrated StarBEAST3 species tree. Tips represent 2 individuals per island. Node bars represent 95% HPD intervals. Branch width denotes effective population size. The top-colored bars represent the lifespan of the extant islands based on island ages from Geist (2014). (B) Plausible paleogeographic reconstruction of the Galápagos adapted from Karnauskas et al. (2018).

#### 3.6 Runs of Homozygosity

Runs of homozygosity (ROH) varied substantially among island populations (Fig. 4A). Española individuals exhibited the highest levels of homozygosity ranging from 181 to 202 ROHs among the two sampled individuals and a cumulative ROH length of 110.0 to 113.9 Mbp, about 61% of the genome. San Cristóbal showed intermediate, though still substantial values, with 68.4 ± 5.0 ROHs and 27.7 ± 1.9 Mb of cumulative ROH per individual. In contrast, Floreana, Isabela, Santa Cruz, and Santiago exhibited markedly lower ROH burden, averaging between 2.1 and 7.2 ROHs per individual and 0.8-2.8 Mb of cumulative ROH. The longest ROH segment observed in Española was 2.34 Mb and San Cristóbal was 1.22 Mb, whereas longest ROH lengths in the remaining populations were all <0.85 Mb.

**Figure 4.**
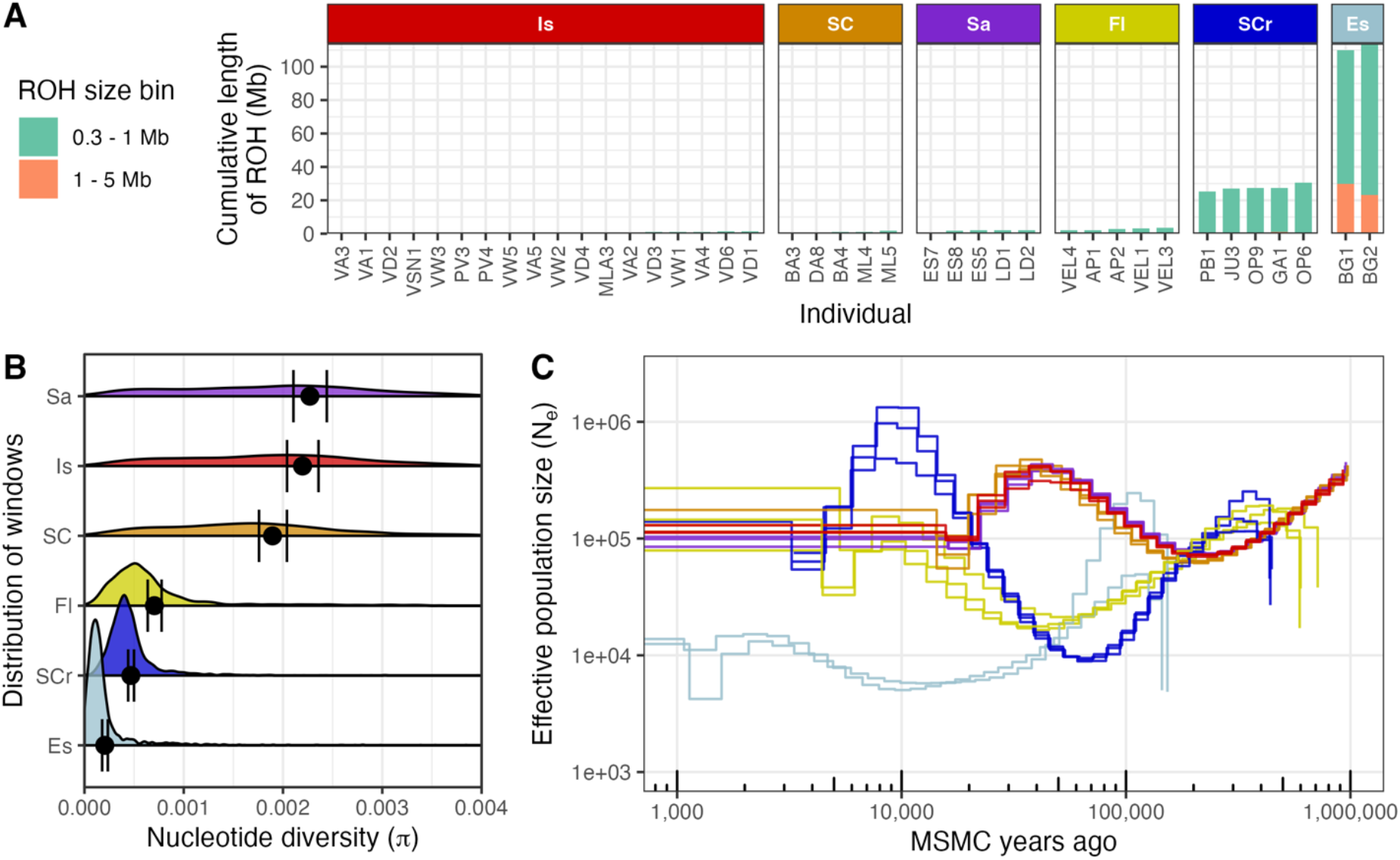
Measures of individual-level inbreeding, population-level genetic diversity, and historical effective population size. (A) Runs of homozygosity (ROHs) per individual. An ROH was defined as any continuous tract of homozygotic sites spanning > 300 kb. (B) Genome-wide distributions of 100 kb windowed π. Points represent the genome-wide mean estimate. Error bars represent 95% confidence intervals obtained from a 1 Mb genomic block bootstraps of 1,000 replicates. (C) Historical demographic histories of the 5 highest coverage individuals per island inferred by MSMC2. The axes were scaled assuming a generation time of 1 year (Gherling et al. 1989) and a per generation mutation rate of 3.5 x 10^-9^ (Liu et al. 2016).

Genome-wide nucleotide diversity (π) varied substantially among populations (Fig. 4B). The highest diversity was observed on Santiago (π = 0.00227) and Isabela (π = 0.00220), followed by Santa Cruz (π = 0.00190). In contrast, lower nucleotide diversity was observed in Floreana (π = 0.000704), San Cristóbal (π = 0.000468), and Española (π = 0.000205). Relative to Española, nucleotide diversity was approximately 11-fold higher on Santiago and Isabela, 9-fold higher on Santa Cruz, 3-fold higher on Floreana, and 2-fold higher on San Cristóbal. Pairwise comparisons of genome-wide nucleotide diversity (Δπ) supported differences among all population pairs, with 95% bootstrap confidence intervals excluding zero for every comparison (fig. S10).

#### 3.7 Historical Demography

Historical demographic histories inferred with MSMC2 were congruent among the central islands of Isabela, Santa Cruz, and Santiago, suggesting that these populations have been shaped by similar historical events (Fig. 4C). In contrast, Floreana, Española, and San Cristóbal each exhibited distinct histories, both relative to the central islands and to one another. Following divergence of all demographic trajectories approximately 100,000-200,000 MSMC years ago, the central island populations showed a gradual increase in effective population size (N_e_), whereas Floreana exhibited a gradual decline, and San Cristóbal a more pronounced reduction in N_e_. Both Floreana and San Cristóbal subsequently showed evidence of demographic recovery toward the present. Following the separation of the Española trajectory from that of San Cristóbal, Española experienced a marked decline in N_e_ that was followed by a prolonged period of relatively low effective population size, with only modest evidence of recent population growth. All populations exhibited relatively stable N_e_ over the past 10,000 MSMC years. However, because our MSMC2 parameterization intentionally merged the most recent time intervals, and because recent estimates are additionally sensitive to sequencing depth, phasing accuracy, and limited sample size, we refrain from drawing strong inferences regarding the most recent demographic history.

#### 3.8 Phenotypic Differences – Body Mass

Body mass varied significantly among islands (likelihood ratio test: *χ*^2^_5_ = 16.65, *p* = 0.0052). Mean body mass ranged from 468 mg on Santiago to 625 and 686 mg on San Cristóbal and Española, respectively, with intermediate values on Isabela (479 mg), Santa Cruz (491 mg), and Floreana (516 mg) (Fig. 5). Pairwise comparisons revealed that bees from San Cristóbal were significantly larger than those from Isabela (*p* = 0.006), Santiago (*p* = 0.020), and Santa Cruz (*p* = 0.024). Differences between San Cristóbal and Floreana were marginally significant (*p* = 0.096). No other pairwise comparisons were statistically significant. Bees from Española tended to be larger than those from the western/central islands, however, estimates for Española were associated with high uncertainty due to limited sample size (n = 4). It should be noted that island-level differences are confounded with non-overlapping sampling periods across islands, and observed differences could therefore be influenced by seasonal environmental variability. However, for the subset of comparisons where temporal sampling can be controlled, we recover the same pattern suggested by the full model. For example, both Santa Cruz and San Cristóbal were sampled in May 2022. Within this subset, bees from Santa Cruz weighed on average 139 mg less than those from San Cristóbal (*t*_33_ = -3.25, *p* = 0.002), consistent with the larger body size observed in San Cristóbal in the full dataset. This result suggests that the observed island-level differences are not solely an artifact of temporal sampling variation.

**Figure 5.**
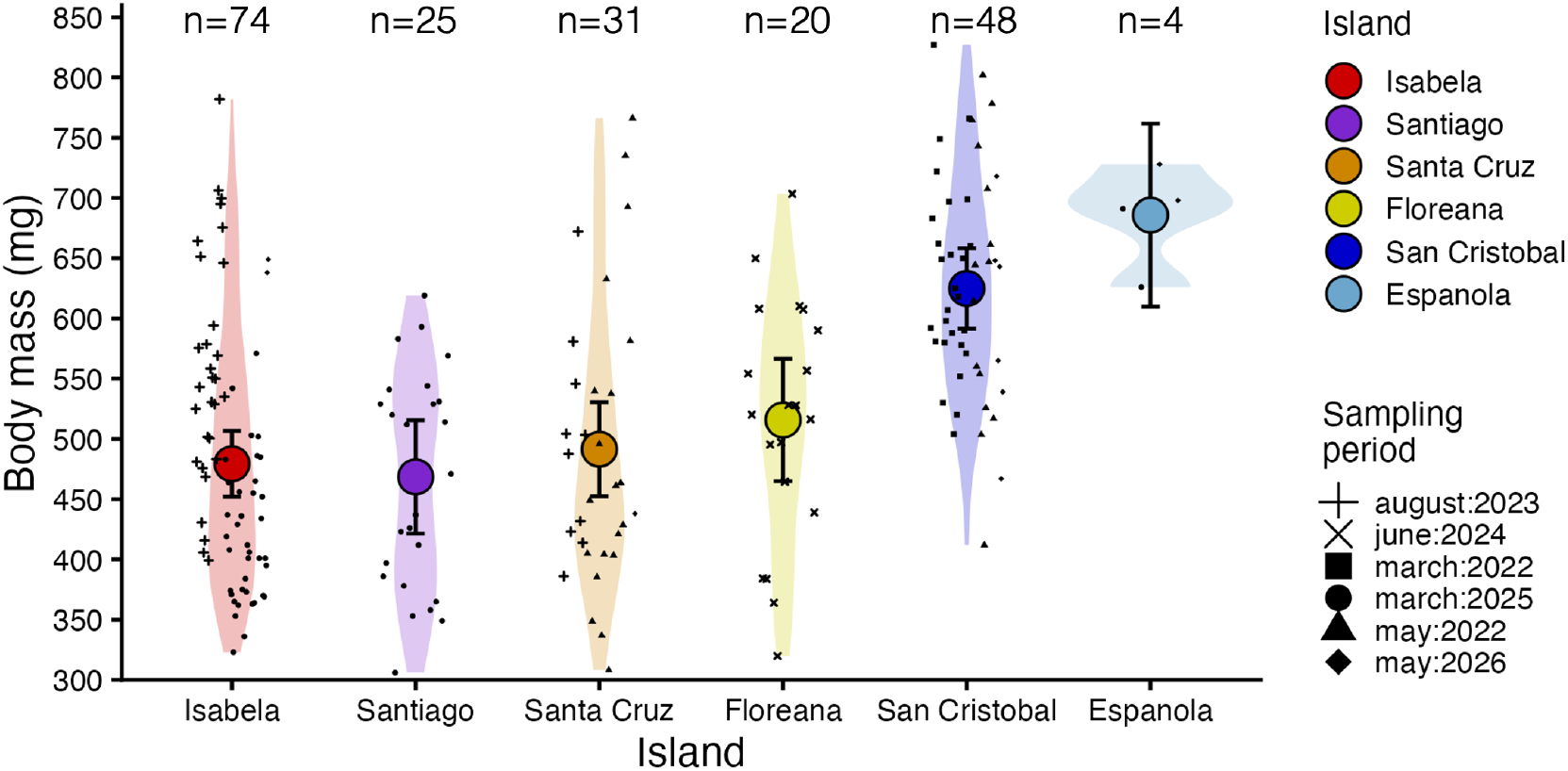
Whole body mass of 202 female *X. darwini* individuals sampled from across 6 islands between 2022 and 2026.

#### 3.9 Expanded UCE Phylogeny

In the expanded UCE dataset containing 3 mainland relatives, the multi-species coalescent (MSC) ASTRAL approach (fig. S11A) and the concatenated approach (fig. S11B) recovered several well-supported terminal clades, placing each individual sampled from the same island as reciprocally monophyletic. In addition, the concatenated analysis strongly supported the expected sister relationship between *X. andica* and *X. transitoria*. In contrast, the ASTRAL species tree recovered *X. andica and X. tranisitoria* along successive branches subtending the group; however, support for these deeper relationships was low to moderate according to posterior probability estimates. Nodes defining the deeper backbone in the ASTRAL tree showed reduced support (*e.g.* local posterior probabilities <0.5 for several internal nodes), indicating uncertainty in the branching order among major lineages. By comparison, the concatenated analysis produced a fully resolved topology with high bootstrap support across most internal nodes, including those defining the relationship between *X. andica*, *X. transitoria* and the rest of the GA samples. Because of the discordance among methods and the limited phylogenetic signal, the UCE dataset does not clearly resolve the relationships between the mainland relatives and the Galápagos lineages, leaving open the possibility that *X. darwini* colonized the Galápagos through multiple independent colonization events.

## 4. Discussion

### 4.1 Population Structure, Island Relationships, and Body Size Differences

Patterns of population structure across the archipelago reveal moderate to strong genetic differentiation among all island populations of the Galápagos carpenter bee, with no evidence of recent or ongoing gene flow between any of the sampled island pairs. This suggests that ocean barriers are effective at preventing recurrent inter-island migration, despite the presumed strong flight ability of the bee (Linsley 1966; Cockerell 1935), echoing conclusions from Vargas et al. (2015). An important exception to this pattern comes from the populations sampled from four volcanoes of Isabela – volcán Sierra Negra, Alcedo, Darwin, and Wolf – which were recovered as a single panmictic population in all analyses. This suggests that the large lava flats separating these volcanoes are not effective barriers to dispersal in this group unlike what has been observed in other taxa (Arteaga et al. 2019; Pozo et al. 2025; Gentile et al. 2009; Gaughran et al. 2024). Interestingly, the eastern islands of San Cristóbal and Española displayed the lowest (though still substantial) relative genetic differentiation, clustered tightly in PCA space, and exhibited relatively shallow absolute divergence, despite being the oldest islands.

The six populations are grouped into two major clades, an eastern clade consisting of San Cristóbal and Española, and a western/central clade consisting of Isabela, Santa Cruz, Santiago and Floreana. The deep genetic separation between these two clades is accompanied by large differences in body size. Individuals from San Cristóbal are about 28% larger (128.5mg) than individuals from the central islands of Santiago, Santa Cruz, and Isabela. Española shows the same pattern; however, the limited number of sampled individuals from this island prohibits definitive conclusions about whether body size is generally larger in the eastern islands than in the western islands. This difference in body size is inversely related to genetic diversity, with the larger-bodied populations having significantly reduced genetic diversity relative to the smaller-bodied island populations. Whether this pattern reflects local adaptation, genetic drift, or both remains an open question. Subtle differences in wing coloration between the eastern and western clades have been suggested by early taxonomic studies (Cockerell 1935) and are consistent with the observed differences in body size, though more work on the morphological differences between these two clades is warranted.

Our findings here are consistent with the ancient east-to-west partitioning of genetic variation observed from *mt*DNA (Vargas et al. 2015), though the relationships within each cluster differ from the *mt*DNA tree of Vargas (2015). The *mt*DNA tree places Floreana as the most derived of the Santa Cruz, Isabela, Santiago, and Floreana quartet whereas our whole-genome phylogeny places Floreana as the most basal of the quartet. This mitonuclear discordance aligns with the high discordance among the windowed nuclear DNA gene trees and is suggestive of incomplete lineage sorting or introgressive hybridization.

### 4.2 Timing of Arrival to the Archipelago

Assuming a single colonization of the archipelago and that our sampling includes the oldest split within the archipelago, our estimates place the minimum time of arrival of *X. darwini* at 1.2-1.6 Mya. This arrival time places *X. darwini* into the middle ground of colonization times of GA fauna and flora. Some lineages, including iguanas and geckos, have been proposed to have colonized the archipelago on proto-landmasses existing before the emergence of the current-day islands on the order of 9-10 Mya (Iguanas, Paradiso et al. 2025) or when only the oldest extant islands were beginning to emerge around 4 Mya (Geckos, Myers et al. 2025). Other lineages began to diversify much more recently – within the last 600 Kya in the case of the yellow warbler (Chaves et al. 2012), or the *Scalesia* giant daisies (Fernandez-Palacios et al. 2020) and *Opuntia* cactuses (Zapata et al. 2024).

Our estimate of a minimum arrival window of 1.2-1.6 Mya should be considered a conservative upper estimate. Because phylogenetic macroevolutionary substitution rates are lower than recent pedigree-based mutation rates, divergence times inferred using these rates may overestimate the age of recent population splits due to the purifying selection and rate averaging over deep timescales (Ho et al. 2011; Lisle and Svensson 2026). Therefore, slightly younger divergence times are possible. Moreover, we can only confidently constrain the timing of colonization between the stem and crown ages of the Galápagos clade. The stem age, defined by the divergence between *X. darwini* and its continental sister taxa, provides the earliest possible time at which colonization could have occurred, whereas the crown age of the Galápagos clade provides the latest possible time by which colonization must have occurred because diversification within the archipelago was already underway. Recent global UCE *Xylocopa* phylogenies which incorporate fossil calibrations from outside Xylocopinae estimate the separation of *X. darwini* from the MRCA mainland species to have occurred somewhere between 1 and 4 Mya (Melo and Martins 2025; Blaimer et al. 2018). A maximum arrival time of 4 Mya would rule out an ancient colonization of the archipelago before the emergence of the oldest current day islands by *X. darwini*. Incorporation of WGS from continental sister taxa would allow for a more precise estimate of maximum arrival time.

Although relatively uncommon, independent colonizations of the islands have occurred in several GA clades including *Phyllodactylus* leaf-toed geckos (Torres-Carvajal *et al*. 2016) and *Hogna* wolf spiders (De Busschere *et al*. 2010). Analysis of mined UCEs from our genome-wide data alongside a broader sample of UCE data including the closest mainland species (Blaimer et al. 2016; 2018) could not rule out this possibility in *X. darwini* (fig. S11). Concatenated and species-tree approaches produced discordant relationships among mainland and island lineages, with the concatenated topology providing some support for two independent colonization events, whereas the species-tree topology supported a single colonization event. However, low support for deeper relationships and disagreement among methods indicate that the number and timing of colonization events remain unresolved. This uncertainty likely reflects the difficulty of resolving rapid divergences using UCE data, where short internal branches and incomplete lineage sorting can obscure phylogenetic signal. Whole-genome data from mainland sister species will be necessary to distinguish among competing colonization scenarios and better resolve the evolutionary history of *X. darwini*.

What predictable factors, if any, led to a successful colonization of *X. darwini* is not currently known. Because of the ecological requirements of carpenter bees in terms of food (nectar) and nesting substrate, the establishment of flowering plants and especially woody plants might have been a necessary prerequisite for the establishment of *X. darwini* (Linsley et al. 1966). Increased phylogenomic studies into the arrival time of endemic plants would help address this question. Interestingly, current estimated arrival times of *Scalesia* giant daisy trees, which *X. darwini* has been found to nest in, are much more recent than our estimate of *X. darwini*. *X. darwini* has been known to nest in the Palo Santo tree (*Bursera graveolens*) and the Galápagos Croton shrub (*Croton scouleri*) which have not yet been studied in a phylogenetic context (Linsley et al. 1966).

### 4.3 Intra-archipelago Diversification and Biogeography

Because this intermediate arrival time occurred after most major islands were already emergent, inferring which island was colonized first is less straightforward than for lineages that arrived when only the oldest islands were subaerial. In those cases, the classical progression rule of island biogeography can be invoked by linking colonization sequence to age of island emergence (Funk and Wagner 1999; Shaw and Gillespie 2016; Rosindell and Phillimore 2011). Here, which island was colonized first might be primarily stochastic or could be driven by more predictable processes (Cowie 2006) such as island size (*e.g.* the target-area hypothesis, Lomolino 1990), ecological filtering (Gillespie and Roderick 2002), competition (Al-Namazi and Bonser 2022), or ocean or prevailing wind patterns (Peck 1991). However, paleo-reconstructions of the geography of the GA during this time period paint the archipelago as consisting of a large central landmass consisting of Santa Cruz, Santiago, and potentially Floreana, surrounded by peripheral islands including Española (Karnauskas et al. 2018; Geist et al. 2014), and other now sunken islands (*e.g.* Sinton et al. 2017), reducing the number of possible landing sites. There is disagreement among paleo-reconstructions of the placement of San Cristóbal with some placing it as isolated for much of its history (Geist et al. 2014), and others depicting it as contiguous with the large central landmass until 1 to 1.5 Mya (Karnauskas et al. 2018). Our estimated divergence time of the eastern and western clade more closely align with the estimated time of vicariant separation suggested by the reconstructions of Karnauskas et al. (2018). A more ancient separation of San Cristóbal from a large central island landmass followed by long-term isolation of San Cristóbal is also consistent with the higher genetic diversity on the central islands compared to the east.

Following the initial separation of the central and eastern Galápagos clades, the central clade underwent a successive series of divergence events beginning with the separation of Floreana (∼698 Kya), followed by Isabela (∼460 Kya), and finally the split between Santiago and Santa Cruz (∼389 Kya). Notably, this sequence does not exactly correspond to the age of emergence of the islands. Instead, diversification appears to follow the order of fragmentation of the formerly united central paleo-landmass reconstructed from paleogeographic models (Geist et al. 2014; Karnauskas et al. 2018; Fig. 3B), suggesting that vicariant fragmentation of this ancestral landmass, rather than sequential colonization of newly emergent islands, drove diversification within the central clade. In contrast, the recent divergence between Española and San Cristóbal (∼61 Kya), coupled with paleogeographic evidence that these islands have not been connected to one another in the recent past (Karnauskas et al. 2018; Ali & Aitchison 2014; Schwartz et al. 2018), strongly supports a founder event from San Cristóbal to Española. Because Española is the oldest island in the archipelago and would be expected to have been colonized much earlier, this recent divergence instead suggests local extinction followed by recolonization, as predicted for islands approaching the end of their geological lifespan (Valente *et al*. 2014; MacArthur and Wilson 1967; Whittaker et al. 2017).

Although fragmentation of the central landmass initiated divergence among the central island populations, climatological modelling suggests this isolation was not permanent. Recurrent reconnections between islands via sea-level-driven land bridges during the late Pleistocene may have provided opportunities for secondary contact following the initial divergence. Models of sea-level change and dating of wave-cut rounded cobbles show that at conservative sea level lowstands, the central islands were reconnected, potentially within the last 20 Kya (Ali & Aitchison, 2014; Schwartz et al. 2018). In contrast, the eastern islands remained isolated throughout these sea-level fluctuations.

These contrasting histories of connectivity are reflected in both historical demographic inference and contemporary patterns of genetic diversity. Historical demographic inference using the sequentially Markovian coalescent (SMC) reconstructs relatively stable and similar effective population histories for Isabela, Santiago, and Santa Cruz, including a notable increase in effective population size between approximately 100,000 and 200,000 MSMC years ago. Floreana, however, shows a gradual decline in effective population size at the same time the gradual increase in effective population size is observed on the central islands, suggesting reduced opportunities for secondary contact compared with the other central islands. The eastern lineage exhibits a markedly different demographic signature. Low nucleotide diversity together with extensive runs of homozygosity indicate long-term isolation with a small effective population size, consistent with the eastern islands having remained disconnected from the central Galapagos during sea-level lowstands (Ali and Aitchison, 2014; Schwartz et al. 2018).

Interestingly, these results are contrary to the expected positive relationship between island area and genetic diversity (Fan et al. 2025). Genetic diversity appears to correspond more closely to island age and historic connectivity, suggesting that island geological history has exerted a stronger influence on long-term population persistence than island size alone. Floreana is broadly consistent with this interpretation. Floreana has ∼3.23 times less land area than San Cristóbal (Snell et al. 1999) yet appears to have maintained a similar historical effective population size and has retained 1.5-fold higher nucleotide diversity than San Cristóbal. Despite Floreana’s earlier separation from central archipelago, its proximity probably permitted prolonged genetic connectivity during the early stages of isolation, potentially providing an example of diversification with a gradual rather than immediate cessation of gene flow. Nucleotide diversity is similarly high across Isabela, Santiago, and Santa Cruz, consistent with more recent gradual vicariant separation of a large, interconnected ancestral population combined with subsequent recurrent interisland reconnections. Exceptionally, Floreana exhibits substantially lower nucleotide diversity than the other central islands, consistent with its earlier separation from the ancestral central landmass. Sequencing of individuals from other smaller islands known to have been contiguous with the ancient central Galápagos landmass such as Pinzón or Santa Fé would lend insight into the relative roles of historical connectivity and island size in shaping patterns of genetic diversity. Although much of the observed demographic signal appears to reflect long-term geological history, more recent demographic changes may also have contributed to the observed patterns of ROH on Española and San Cristóbal. For example, Holocene lava flows may have substantially reduced suitable habitat and fragmented populations on San Cristóbal (McLeod et al. 2015).

## Conclusion

Our study highlights the complexities that shape population history and contemporary genetic variation in a highly dynamic insular system. We find strong genetic isolation among island populations and no evidence of contemporary gene flow between sampled island pairs suggesting that successful overwater dispersal of *X. darwini* appears to be a rare occurrence. All island pairs except for the two eastern islands show relatively high pairwise genetic divergence and split times occurring on the order of hundreds of thousands of years ago. Under a single-colonization scenario, almost all major extant islands (except for Isabela and Fernandina) were already subaerial and available for colonization (Geist et al. 2014). Between approximately 0.5 and 1.5 Mya a large paleo-landmass underwent a series of fragmentation events driven by tectonic and volcanic activity. Rather than mirroring island age, the diversification closely follows the hypothesized sequence of island fragmentation beginning with the separation of San Cristobal followed by the central and western islands of Floreana, and then Isabela, Santiago, and Santa Cruz (Karnauskas et al. 2018). This pattern is consistent with vicariance being the primary driver of diversification in this lineage and highlights that simple predictions like the progression rule cannot always be invoked in geologically dynamic systems like the Galápagos. We interpret the shallow split between the eastern islands of Española and San Cristóbal as consistent with a recent founder dispersal event and indicative of possible local extinction-recolonization dynamics on older islands. Notably, this is the only example of over-water founder dispersal explaining diversification among the sampled islands. Measures of contemporary genetic diversity and runs of homozygosity (ROHs) combined with reduced historical effective population sizes on the southern island of Floreana and especially the eastern islands of Española and San Cristóbal are consistent with the long-term isolation of these islands, even during sea-level lowstands. These measures of diversity do not correlate with island size but instead likely reflect a combination of historical demographic events such as population contraction driven by island fragmentation, founder effects following dispersal-driven colonization, island erosion and subsidence, and eustatic inter-island reconnections. Our results suggest that the Galápagos carpenter bee is likely not a frequent overwater disperser, despite the historical presumption of strong flight ability. The resulting lack of recurrent interisland migration has allowed historical, landscape-driven signatures of diversification to remain preserved within the genome.

## Funding Statement

This work was supported by the U.S. National Science Foundation under award number 2212157 to BH, a University of South Carolina SPARC grant to GCM, USFQ Galápagos Grants awarded to MLT in 2023, 2024, and 2025, and to MJP in 2026, as well as a USFQ COCIBA Grant awarded to MJP in 2023.

## Data Statement

All scripts used in the analyses can be found at https://github.com/graham-evo/Xdarwini.Manuscript.git. Trimmed short reads and HiFi long reads are deposited in NCBI under PRJNA1448021. Other data including R scripts to produce figures can be found on Dryad.

## Acknowledgements

We would like to thank all members of the Galápagos Science Center (GSC) who contributed to the execution of this project, especially Paúl Yepez for logistical support. We are also grateful to the members of the Plant Biotechnology Laboratory for their support, especially Milton Gordillo and Daniel Dávila. We also would like to thank students and colleagues from the University of South Carolina who aided in bee collections, including Celia Razon, Naomi Wehmeir, Victoria Guy, David Conroy, Joel Cheek, Allison Kalvitz, Emma Harrison, Thelma Zaw, Kate Gahlhoff, and Kevin Ayres. We thank Carrie Wessinger and Nitin Ravikanthachari for helpful discussions, Ben Stone for assistance with bioinformatics tools, and Ricky Flamio for advice on genome assembly. The authors gratefully acknowledge the computational resources provided by the Hyperion high performance computing cluster at the University of South Carolina. We also acknowledge the technical assistance and resources provided by Research Computing at the University of South Carolina (RRID:SCR_027488). Finally, we thank the Galápagos National Park Directorate for authorizing the research conducted under permits MAATE-DBI-CM-2022-0268 and MAATE-DBI-CM-2025-0460.

## Author Contributions: GCM

Methodology, Formal Analysis, Investigation, Writing - Original Draft, Visualization **VBM:** Formal Analysis, Investigation, Visualization, Writing - Review & Editing **MJP:** Investigation, Project administration, Writing - Review & Editing **MLT:** Conceptualization, Investigation, Resources, Writing - Review & Editing, Supervision, Project administration, Funding acquisition **BH:** Conceptualization, Methodology, Investigation, Resources, Writing - Review and Editing, Supervision, Funding acquisition.

## 2. Materials and Methods

### 2.1. Sampling, DNA Extractions, and Sequencing

We sequenced 42 diploid female individuals and a single haploid male individual (for *de novo* genome assembly) of *Xylocopa darwini* collected across 6 islands of the Galápagos archipelago (GA) on sampling trips conducted in 2023, 2024, and 2025 (Fig. 1). On Isabela, we collected individuals from four volcanoes (Cerro Azul, Sierra Negra, Alcedo, and Wolf), which existed as separate islands until approximately 800 Kya before merging into their present-day configuration (Harpp and Geist 2018). Individuals were collected using a butterfly net and were stored on ice until DNA extraction. See supplemental material S1 (Excel Sheet 1) for sampling details.

For whole-genome resequencing of diploid females, DNA was isolated from 30 mg of dissected abdominal tissue using the DNeasy Blood and Tissue Kit (Qiagen) following manufacturer protocols. DNA purity was assessed on a NanoDrop Lite spectrophotometer (ThermoFisher Scientific), fragment length was checked on an agarose gel, and DNA concentration was estimated using a Qubit4 fluorometer with broad range dsDNA working solution (ThermoFisher Scientific, Invitrogen). Illumina 150 bp paired-end short-read sequencing libraries were prepared by tagmentation and sequenced in two independent runs of an Illumina NovaSeqX instrument at the Duke Sequencing and Genomic Technologies Core Facility in November 2023 and August 2025. Reads were demultiplexed into sample-specific FastQ files by the core facility. Reads from separate lanes were concatenated using a custom bash script. See Supplemental Material S2 for extraction and sequencing details (Excel Sheet 2).

### 2.2 *de novo* Draft Genome Assembly

High molecular weight (HMW) DNA was isolated from abdominal tissue of a haploid male collected from San Cristóbal using the E.Z.N.A DNA Extraction Kit (Omega BioTek). DNA quality and concentration were checked with a NanoDrop spectrophotometer, and fragment length was confirmed on an agarose gel. SMRTbell long-read libraries were prepared with the SMRTbell Revio Prep Kit (Pacific Biosciences) and Circular Concensus Sequences (CCS) were generated by single molecular real time (SMRT) sequencing on the Sequel II sequencing platform (PacBio) at the Arizona Genomics Institute (University of Arizona). CCS reads were processed with PacBio’s CCS software (v7.0.0), to produce high fidelity (HiFi) long reads (Wenger et al. 2019) in unaligned binary alignment format (uBAM) before finally being converted to FastQ files with Samtools v1.5 bam2fq. Raw HiFi long reads are available at the NCBI SRA under the BioProject PRJNA1448021.

We initially assembled reads *de novo* into four draft genomes using the HiCanu (v2.2.0) (Nurk et al. 2020), Hifiasm (v0.19.8) (Cheng et al. 2021), Verkko (v1.4.1) (Rautiainen et al. 2023), and HiFlye (v2.9.3) (Kolmogorov et al. 2019) assembly algorithms. QUAST (v5.2.0) (Mikheenko et al. 2018) was used to compute assembly contiguity metrics (N50, L50). Assembly completeness was assessed by the presence of basic universal single copy orthologous genes (BUSCOs) with BUSCO (v5.4.7) (Simão et al. 2015) by searching the hymenoptera_obd10 lineage BUSCO database, which at the time of retrieval (01-13-2024) contained 5991 BUSCOs harvested from 40 hymenopteran genomes. Haploid assembly size was estimated by counting k-mers of k=19 and k=31 with Jellyfish (v3.0) (Marçais and Kingsford 2011) and analyzing the k-mer count distribution with GenomeScope (v1.0) (Vurture et al. 2017). Based on assembly contiguity and completeness, we chose the HiFlye assembly for all downstream analysis. Contigs were further filtered after mapping (see section 2.3).

### 2.3 WGS Data - Read QC, Mapping, Variant Calling, and Datasets

Raw Illumina paired-end reads (150 bp) were trimmed using fastp (v0.23.4) (Chen et al. 2018) to remove adapter sequences, low-quality bases, and poly-G tails. For each sample, forward and reverse reads were processed in paired-end mode and adapter detection was performed automatically (--detect_adapter_for_pe), with Illumina adapter sequences redundantly specified for both read directions. polyG artifact trimming was enabled to remove homopolymer artifacts common to Illumina NovaSeq data. Trimmed reads were written to compressed FastQ files, while per-sample quality reports were generated and aggregated into a single summary report using MultiQC (v1.15) (Ewels et al. 2016). Trimmed reads are available at the NCBI SRA under the BioProject PRJNA1448021.

Forward and reverse quality-trimmed read pairs from the 42 diploid GA samples were mapped to the *X. darwini* HiFlye draft genome assembly using BWA-MEM (v0.7.17) (Li 2013) Read group information (sample ID, library, platform) was added during mapping using Picard. HiFi long read data for the outgroup, *Xylocopa virginica* was retrieved from the NCBI SRA (Accession # SRR31041156) and mapped to the *X. darwini* HiFi draft assembly using Minimap2 (v2.24) (Li 2021) --hifi mode.

Mapped reads were piped through a post-processing pipeline using SAMtools (v1.15) (Danecek et al. 2021) and bamUtil (v1.0.15) (Jun et al. 2015). Reads were first name-sorted and processed with samtools fixmate to ensure mate pair information was correct, followed by coordinate sorting with samtools sort. Duplicate reads were identified and removed with samtools markdup. Properly paired reads with a mapping quality of ≤ 30 were flagged as unmapped but kept in the alignment file. Overlapping bases between mates were soft-clipped using bam clipOverlap to avoid double-counting in downstream depth, altering allele frequency calculations. The resulting Binary Alignment Map (BAM) files were indexed with samtools index and validated with samtools quickcheck. Coverage and mapping statistics were generated with samtools coverage (histogram and tabular outputs) and samtools stats. To provide fine resolution windowed coverage summaries and to inform depth filtering thresholds, MosDepth (v0.3.8) (Pedersen and Quinlan 2017) was run in 1 kb windows and thresholds of 1×, 5×, 10×, 20×, and 30×. Additional quality metrics and graphical reports were produced with QualiMap bamqc (v2.3) (Okonechnikov et al. 2015). All quality control outputs, including SAMtools summaries, QualiMap reports, and MosDepth statistics, were aggregated into a single interactive report using MultiQC.

We evaluated and filtered contigs based on mean mapping quality (MAPQ), breadth of coverage, and median sequencing depth. We applied a stepwise contig filtering process, removing contigs if they did not meet the threshold for each metric in at least 90% of our GA samples (38/42 individuals). We removed contigs with mean MAPQ < 30, <90% breadth of coverage, high read depth (defined as exceeding the 95th percentile of read depth across all mapped sites). This left us with a reference set containing 24 core contigs spanning 185,432,943 base pairs. We performed variant calling on the mapped reads of the 42 GA individuals plus the outgroup for each of the 24 contigs separately using a combination of VCFtools v0.1.17 (Danecek et al. 2011) and BCFtools v1.11 (Danecek et al. 2021).

We generated a number of datasets for downstream analyses (See TableS3 for details). Briefly, we began by generating a quality-filtered SNP dataset (Dataset A) by excluding indels and invariant sites and removing the outgroup. SNPs were retained if they had a minimum quality score of Q30, mean depth between 10× and 40× across samples, per-genotype depth between 10× and 60×, and less than 20% missing data across individuals. This dataset was further refined by excluding low-frequency variants (minor allele frequency <0.02) to produce Dataset B. To generate Dataset C, which included both variant and invariant sites, we concatenated the quality-filtered SNPs (Dataset A) with a 20% missingness filtered set of invariant sites, again excluding the outgroup. Finally, we produced a strictly biallelic SNP Dataset D by restricting Dataset A to sites with exactly two alleles and created a final Dataset E that included only biallelic SNPs with MAF > 0.02.

For phylogenomic analyses, we filtered variants based on the same criteria described above, this time including the outgroup, producing a dataset of polarized phylogenetically informative SNPs (Dataset F). Invariant sites were then extracted from the raw VCF after removing indels and applying a 20% missingness filter, before combining with Dataset F to produce a Dataset G containing high quality polarized variant and invariant positions.

### 2.4 Population Structure of GA samples

To determine if individuals sampled across islands represented genetically distinct populations we conducted several complementary population genetic analyses. We produced a linkage-pruned SNP panel containing 125,664 biallelic SNPs drawn from a larger sample of 1,737,064 SNPs present at a minor allele frequency of >0.02 across all individuals. To prune, PLINK (v1.90) (Purcell et al. 2007) was used to remove SNPs with an *r*^2^ threshold > 0.1 in sliding windows of 50 SNPs with a step size of 10 SNPs. We conducted a principal component analysis using PLINK’s --pca function on this same linkage-pruned dataset.

The quantify the degree of population genetic structure, we used pixy v1.2.4 (Korunes and Samuk 2021). We estimated absolute genetic divergence *d_xy_* and relative genetic differentiation F_ST_ from a VCF file including both variant and invariant sites to properly account for missing data by restricting denominators to callable sites. We computed each statistic in non-overlapping 100 kb windows by grouping individuals by island and in a separate analysis treating each volcano of Isabela as a distinct group. To reduce non-independence of windowed estimates, significant genetic differentiation was assessed by testing if each pairwise-F_ST_ differed reliably and consistently from 0 when averaged across randomly sampled 100 kb windows. We produced an unrooted neighbor-joining tree using pairwise *d_xy_* values computed between individuals to confirm reciprocal monophyly of individuals sampled from different islands.

### 2.5 Species Tree Inference

To estimate a species tree, we began by estimating gene trees from non-overlapping 20 kb windows. Per-sample consensus sequences were generated from the combined SNP and invariant dataset using bcftools consensus, specifying the reference genome as a template. Heterozygous sites were encoded using IUPAC ambiguity codes (--impact-codes) to retain within-individual variation. Sites with missing genotype calls were represented as “N” (--missing N), while positions absent from the VCF were not included in the output. We then split genome-wide consensus sequences into 24 contig-specific FASTA files and aggregated across all individuals to produce per-contig multiple sequence alignments. We then divided each contig alignment into 20 kb non-overlapping alignments using a custom python script adapted from Stone and Wessinger (2024). Windows were excluded if they contained ≥80% missing data (N’s) for any one individual. For each alignment, the best-fit substitution model was selected using ModelFinder (-m MFP) (Kalyaanamoorthy et al. 2017), and maximum-likelihood trees were inferred using IQ-TREE2 v2.3.6 (Minh et al. 2020). All gene trees were rooted using the outgroup *X. virginica.* This process resulted in 8,878 maximum-likelihood gene trees.

Species tree estimation from all gene trees was conducted using ASTRAL-IV v1.22.4.6 (Zhang et al. 2025) with branch lengths computed using integrated CASTLES-II (Tabatabaee et al. 2023). Individual tips were assigned to the 6 island populations. To improve tree search, the default search was extended using the -R option, resulting in 16 rounds of replacement and subsampling. Branch support was assessed using local posterior probabilities by specifying the -u 2 option, allowing support to be calculated for all alternative resolutions of each branch. The tree was rooted with *X. virginica* and the prompt was run with the random seed 2526. The final species tree was selected based on maximum quartet score, which maximizes topology concordance across gene trees.

### 2.6 Divergence Time Estimation

We estimated a time-calibrated species tree using the StarBEAST3 package v1.1.9 (Douglas et al. 2022) implemented in BEAST v2.7.7 (Bouckaert et al. 2019) with guidance from Taming the BEAST (Barido-Sottani et al. 2017). To improve computational efficiency and model fit, analyses were limited to a subset of 20 kb genomic windows exhibiting relatively clock-like substitution patterns, sufficient phylogenetic signal, and concordance with the species tree. Candidate windows were identified using scripts from the SortaDate pipeline (Smith et al. 2018). For each gene tree inferred in Section 2.6, we excluded the outgroup and calculated root-to-tip variance (a proxy for clock-likeness), total tree length (reflecting phylogenetic signal), and a measure of bipartition support that captures concordance with the species tree. The distributions of these metrics were inspected to determine appropriate filtering thresholds. Loci were retained if they exhibited intermediate tree lengths (0.005-0.03 substitutions per site) and low root-to-tip variance (≤1x10^-5^), thereby excluding both low-information and highly rate-heterogeneous windows. The filtered windows were then ranked by bipartition support, followed by tree length, and root-to-tip variance, and the top 100 windows were selected for downstream analysis. For each retained window locus, we down sampled the dataset to include two individuals per island population prior to divergence time estimation in BEAST2. The 100 chosen window alignments were imported into the software *beauti* v2.7.8 to produce the *.xml* file containing all StarBEAST3 model parameters. Each window was allowed an independent substitution model, tree model, and clock model. The species tree topology was fixed during divergence time estimation to the topology recovered by ASTRAL by removing all operators acting on the species tree topology. Since fossil evidence is rare in the Galápagos, we applied a strict species tree clock rate of 8.4 x 10^-4^ substitutions site^-1^ Myr^-1^ following previously published nuclear substitution rates of bees (Ronquist et al. 2012; Almeida et al. 2023; Spasojevic et al. 2020). We used a calibrated Yule species tree prior to incorporate geological constraints, following Phillips et al. (2019), by placing a broad log-normal prior on the crown age of the Galápagos clade, with a mean (in real space) of 4 Mya and a standard deviation of 0.85 which fit a probability distribution with the most likely arrival time centered on the age of the current islands with a tail extending to 20 Mya to reflect possible earlier colonization on the now sunken islands (Christie et al. 1992). We ran four independent replicate runs of 25 million Markov-chain Monte Carlo (MCMC) steps, sampling posterior trees and model parameters every 2500 steps. Chain convergence was assessed by comparing parameter traces among runs, using Tracer v1.7.2 (Rambaut et al. 2018). After confirming effective sample size (ESS) values >200 for all key parameters in each run, MCMC chains were combined across runs, discarding the first 10% of samples of each run as burn-in, using LogCombiner v2.7.7. The maximum clade credibility species tree was estimated using TreeAnnotator v2.7.7. Both tools are members of the BEAST2 family of packages (Bouckaert et al. 2019).

We did not estimate the substitution rate in either of our analyses because substitution rate and divergence time are not independently identifiable in the absence of strong temporal calibrations (Drummond et al. 2006; Ho & Dechêne, 2014). Preliminary analyses allowing the rate to vary under a log-normal clock rate prior with the broad constraint on the crown node of the Galápagos resulted in poor MCMC mixing and non-convergence. We therefore fixed the substitution rate to a biologically reasonable value to ensure model identifiability and stable inference of divergence times.

### 2.7 Demographic History

We used MSMC2 v2.1.4 (Schiffels and Wang 2020) to estimate demographic histories following the recommendations of Mather et al. (2019). For each analysis we restricted analysis to 16 contigs > 1 Mb in length following the simulations of Gower et al. (2018). We began by generating high-quality phased haplotypes for our diploid female *X. darwini* individuals. We isolated strictly biallelic sites and applied a strict filter removing sites with more than 5% missing data (>2 individuals). Haplotype estimation and missing genotype imputation was performed using Beagle v5.4 (Browning et al. 2021; Browning et al. 2018) with default parameters except for the window size which was set to 10 cM. We concurrently generated an unbiased genomic coverage ‘callable’ mask per individual. For each individual sample, the raw unphased all-sites VCF (containing invariant sites, SNPs, and indels) was filtered to convert per-sample low or high coverage positions and indels to missing (./.), defined as 0.5 times or 2 times the mean coverage of that sample. These were then split by chromosome and processed using the vcfAllSiteParser.py script from https://github.com/stschiff/msmc-tools. This tool generates individual mask files. A separate genome-wide mappability mask was produced using the SNPable seqbility toolkit available from https://github.com/lh3/misc). The reference genome was fragmented into all possible k-mers of n=35 with splitfa which were aligned back to the reference with BWA (Li and Durbin 2009). The resulting alignments were processed with gen_raw_mask.pl to create the precursor mappability mask which was converted to another mappability mask using gen_mask with a mapping threshold of 50%. Finally, this mask was converted to the MSMC2 BED-format compatible mask file using the makeMappabilityMask.py from the aforementioned MSMC-tools GitHub repo. Finally, individual-specific coverage masks, global mappability masks, and beagle-phased variants were integrated into cross-population input files using generate_multihetsep.py. Genome-wide data quality was checked using getStats.d which confirmed a final, high-quality intersected dataset of 70.3 Mb across the 16 core contigs. Chromosomes were passed as separate file arguments to MSMC2. Time intervals were defined using the pattern 1*3+30*1+1*4, resulting in 30 free parameters bounded by merged intervals at the youngest and oldest portions of the timescale, thereby concentrating the free parameters across the intermediate portion of the inferred demographic history.

### 2.8 Runs of Homozygosity (ROH) and Nucleotide Diversity (**π)**

We used PLINK to measure runs of homozygosity (ROHs) (Purcell et al. 2007). The same phased per-contig per-individual singleton-free VCFs and callable masks used for MSMC2 were converted to PLINK compatible .bed .bim and .fam file formats. We then used plink merge-list to produce a master population file containing SNPs for all 42 individuals. Runs of homozygosity (ROH) were identified using PLINK’s --homozyg function. ROHs were defined as homozygous segments spanning at least 300 kb (--homozyg-kb 300) and containing a minimum of 50 SNPs (--homozyg-snp 50). A sliding window of 50 SNPs was used to evaluate homozygosity (--homozyg-window-snp 50) allowing up to two heterozygous genotypes (--homozyg-window-het 2) and five missing genotypes (--homozyg-window-missing 5) per window to account for genotyping error and missing data. Adjacent homozygous segments separated by gaps of ≤ 1Mb were permitted (--homozyg-gap 1000).

Genome-wide nucleotide diversity (π) was estimated from the pixy 100 kb non-overlapping averages (See Section 2.5) using the fraction of summed pairwise differences to summed pairwise comparisons across windows following the recommendations of Samuk et al. (2021). To account for non-independence among neighboring genomic regions, uncertainty in genome-wide π was assessed using a genomic block bootstrap approach. Windows were assigned to contiguous non-overlapping 1 Mb blocks. For each bootstrap replicate, genomic blocks were sampled with replacement, with the number of sampled blocks equal to the total number of observed blocks in the genome. A total of 1,000 bootstrap replicates were performed. Pairwise differences in nucleotide diversity were calculated among populations for each bootstrap replicate, and statistical support was assessed by whether the 95% bootstrap confidence intervals of the pairwise difference excluded zero.

### 2.9 Phenotypic Differences – Body Mass Measures

Whole body mass of the 42 sequenced females and an additional 160 frozen specimens of female *X. darwini* collected from 2022 to 2026 was measured on a microbalance (Metler Toledo). To test for differences in body mass between islands we fit a linear mixed-effects model using the mixed() function implemented in the afex package v2.0.1 (Singmann et al. 2025), a wrapper for lme4 (Bates et al. 2015), in R v4.5.2 (R Core Team 2025). To account for spatial structure of sampling, locality was included as a random effect nested within each island. Because sampling was conducted across multiple years and not all localities were sampled evenly through time, temporal variation in body size could not be accounted for in the model. Model significance for fixed effects was evaluated using likelihood-ratio tests. Estimated marginal means for each island and pairwise comparisons among islands were calculated using the emmeans package v2.0.3 (Lenth and Piaskowski 2017).

### 2.10 UCE Tree Incorporating Mainland Relatives

To establish whether the Galápagos *X. darwini* populations originated from a single or multiple independent colonizations, we tested for monophyly of all island taxa by inferring a phylogenetic tree using published ultraconserved element (UCE) data containing both of the known closest related mainland sister taxa of *X. darwini*: *X. andica* and *X. transitoria* (Blaimer et al. 2018; Blaimer et al. 2016), and an additional outgroup, *X. varipuncta*, estimated to be around 10 million years diverged from the aforementioned clade (Blaimer et al. 2018). To match the completeness of the available UCE data of the outgroups, we reduced our demultiplexed, quality trimmed and adapter-free WGS short-reads of a randomly selected subset of 12 *X. darwini* individuals (2 per island sampled) to UCE-only FASTA files. Reference-guided alignment of our reads to Faircloth (2014)’s Hymenoptera UCE probe set was conducted using aTRAM v2.4.4 (Allen et al. 2018) by building an aTRAM database from the 12 WGS libraries with *atram_preprocessor.py*. We then assembled contigs of these 12 WGS libraries and the 3 mainland sister taxa using the Trinity wrapper within aTRAM’s *atram.py* script. The PHYLUCE package v.1.7.32 (Faircloth et al. 2012; Faircloth 2015) was used to further process the UCE libraries. Sample-specific contig assemblies were matched to specific UCEs within the Hymenoptera UCE probe set using *match_contigs_to_probes.py* (min_coverage = 80, min_identity = 80). *phyluce_assembly_get_match_counts.py* was used to query the relational database containing matched probes created in the previous step to generate a list of UCE loci shared across all samples. We then created FASTA files from this list of UCE loci containing sequence data for all samples sharing that UCE. We calculated the mean length of the UCE assemblies per sample using *phyluce_assembly_explode_get_fastas_file.py* and *phyluce_assembly_get_fasta_lengths.py* finding that the WGS mined UCEs were around 1.6 kbp longer than the UCE-capture samples mostly containing flanking regions not enriched in the UCE capture probes Blaimer et al. (2016). We therefore opted to aggressively trim these flanking regions to reduce the amount of missing data between the WGS samples and the UCE-capture samples. FASTA files for each UCE locus were aligned using MAFFT implemented in *phyluce_align_seqcap_align.py* and internally trimmed using Gblocks (Talavera and Castresana 2007) implemented in *phyluce_align_get_trimmed_alignments_from_untrimmed*.py specifying that at least 90% of samples contain no missing data at that site by calling --proportion 0.9. Finally, we produced a 75% complete data matrix using *phyluce_align_get_only_loci_with_min_taxa.py* and converted this matrix into NEXUS format using *phyluce_align_concatenate_alignments.py*. Relationships of the *X. darwini* island samples to the mainland sister species were inferred from a 75% complete concatenated alignment of 440 UCE loci (238,047 bp) using a maximum likelihood approach using IQ-TREE v2.3.6 (Bui Quang Minh et al. 2020). Model selection was performed using ModelFinder (Kalyaanamoorthy et al. 2017) and a maximum likelihood tree was inferred under the best-fitting TVM+F+I+G4 model, with support estimated using 1,000 ultrafast bootstrap replicates with UFBoot2 (Diep Thi Hoang et al. 2018). We also estimated a MSC species tree with ASTRAL-IV by producing gene trees for each of the 440 UCE loci, again with 1,000 ultrafast bootstrap replicates. For each tree, we collapsed internal branches with BS support values <=10 as recommended by Mirarab and Warnow (2015).

## Supplemental Material

**Figure S1.**
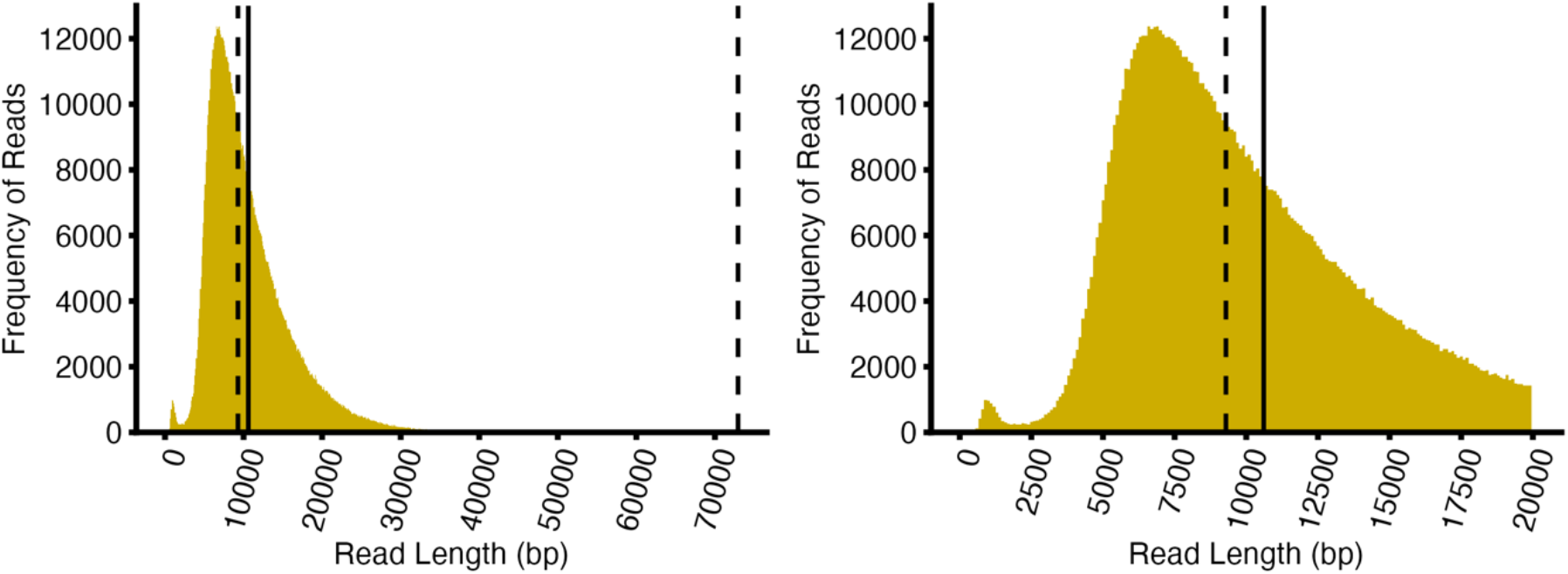
Read length distribution of PacBio HiFi long reads. (Left) Distribution of read length showing the largest reads. (Right) Distribution of read lengths zooming in on bulk of distribution. Dotted line is the median read length; solid line is mean read length.

**Figure S2.**
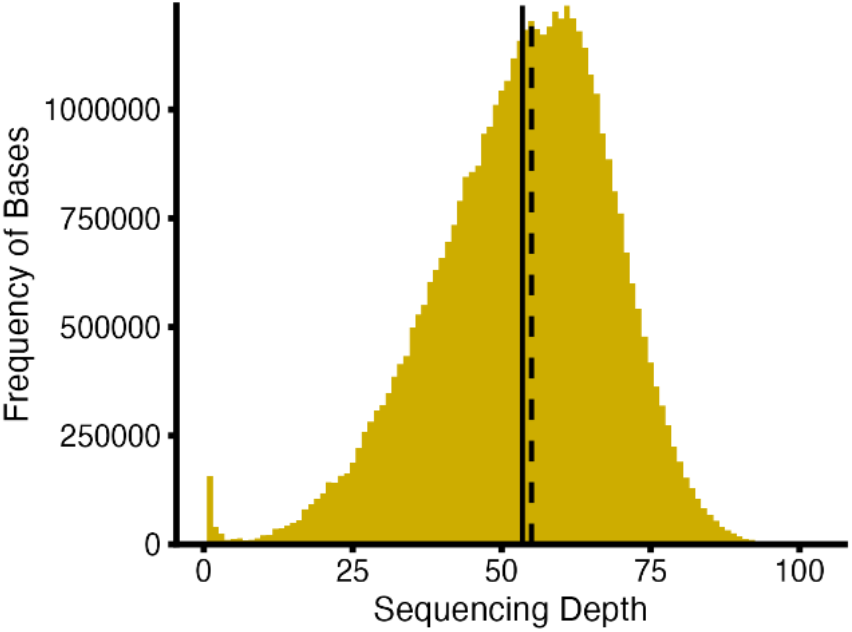
Depth of coverage distribution of PacBio HiFi long reads. Dotted line is the median read length; solid line is mean read length.

**Figure S3.** Draft genome summary statistics displayed using SnailPlot in BlobToolkit (Challis et al. 2020).

**Figure S4.**
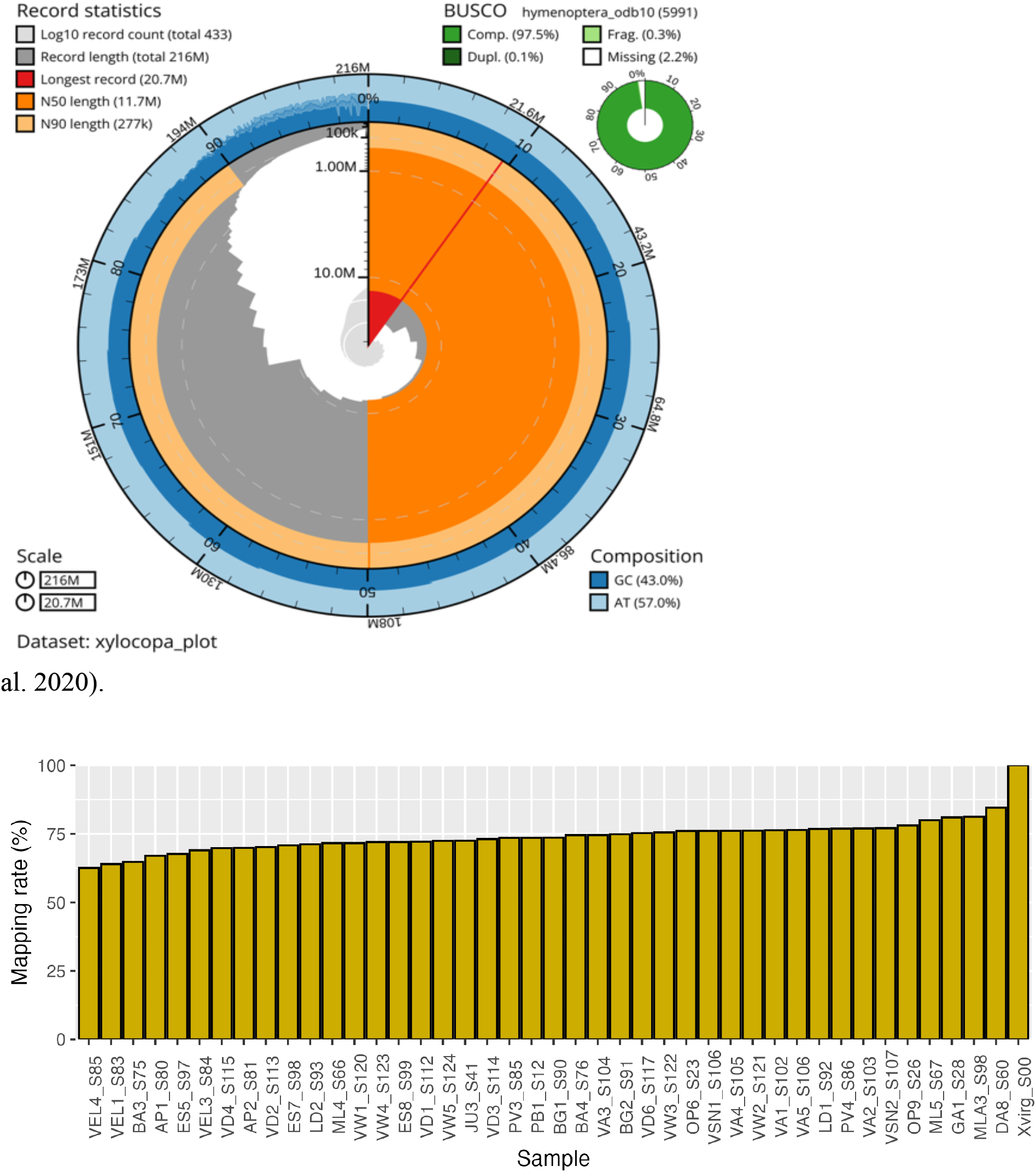
Percent reads mapped per sample.

**Figure S5.**
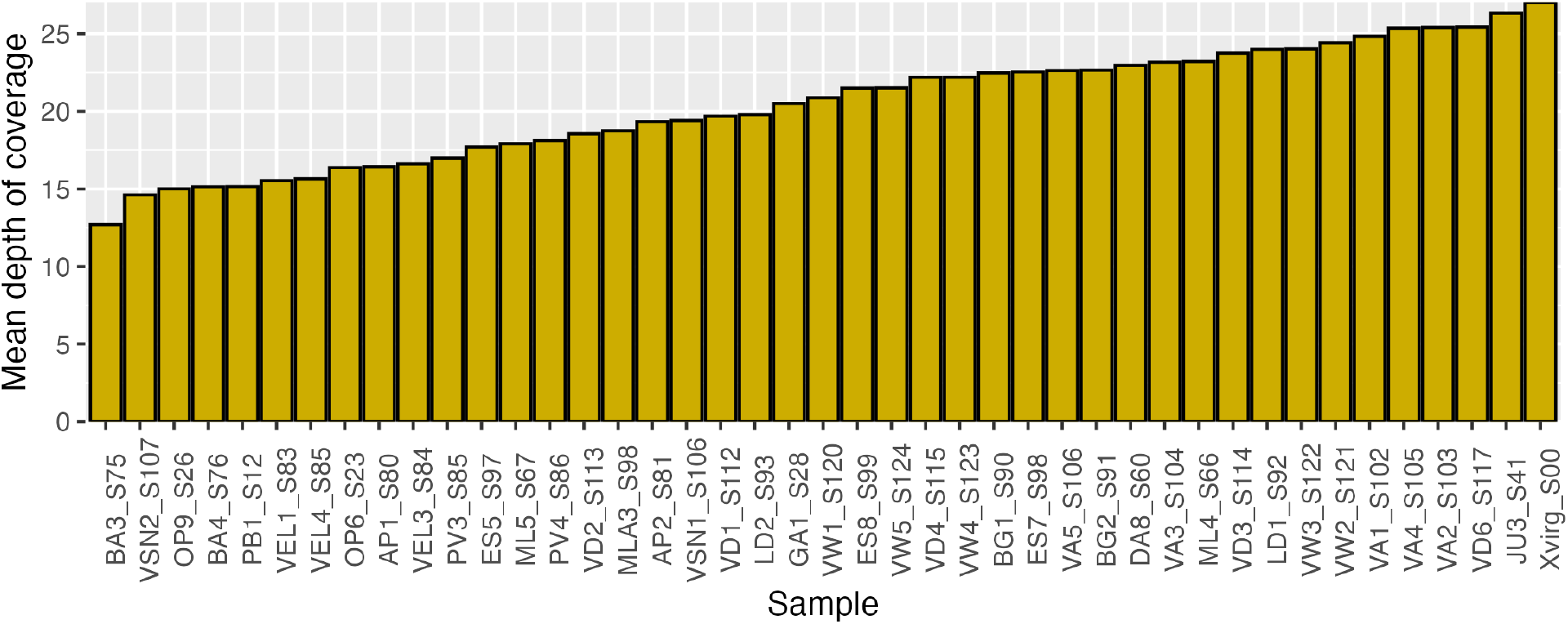
Mean depth of coverage per sample before variant filtering.

**Figure S6.**
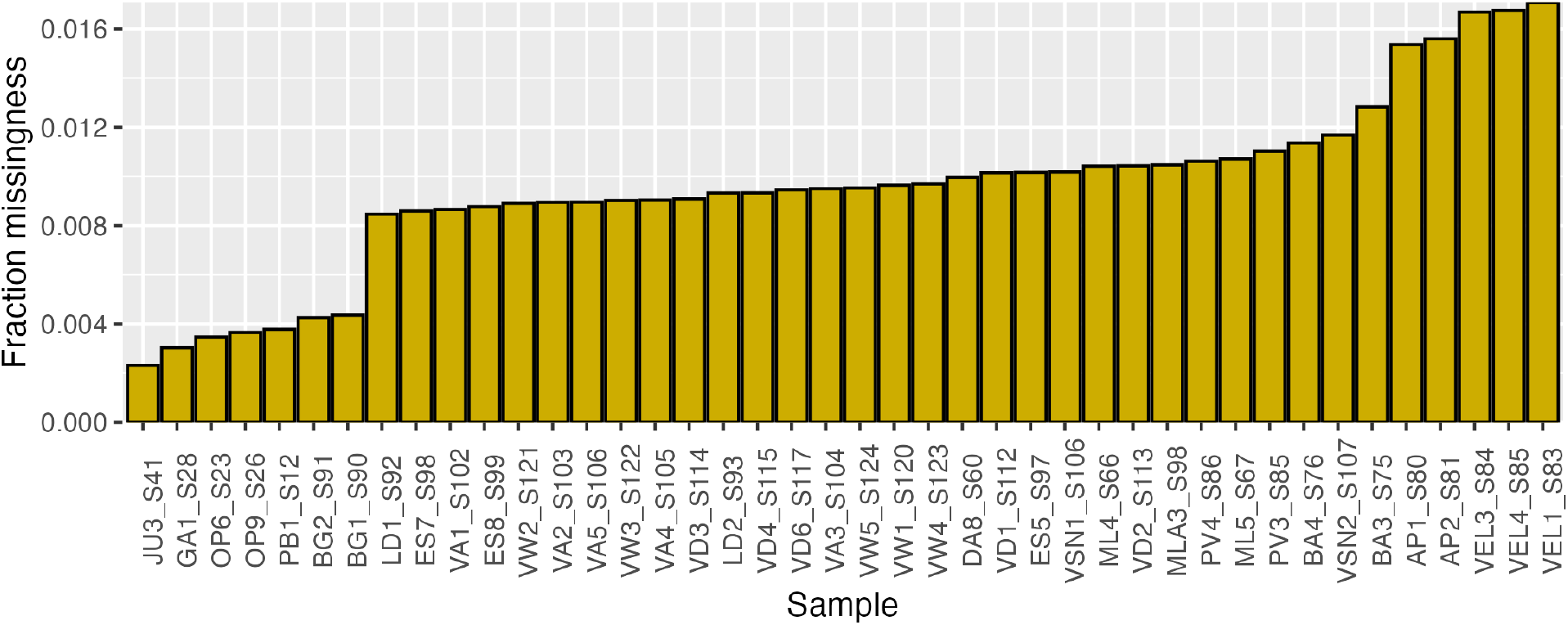
Fraction missing per sample.

**Figure S6.**
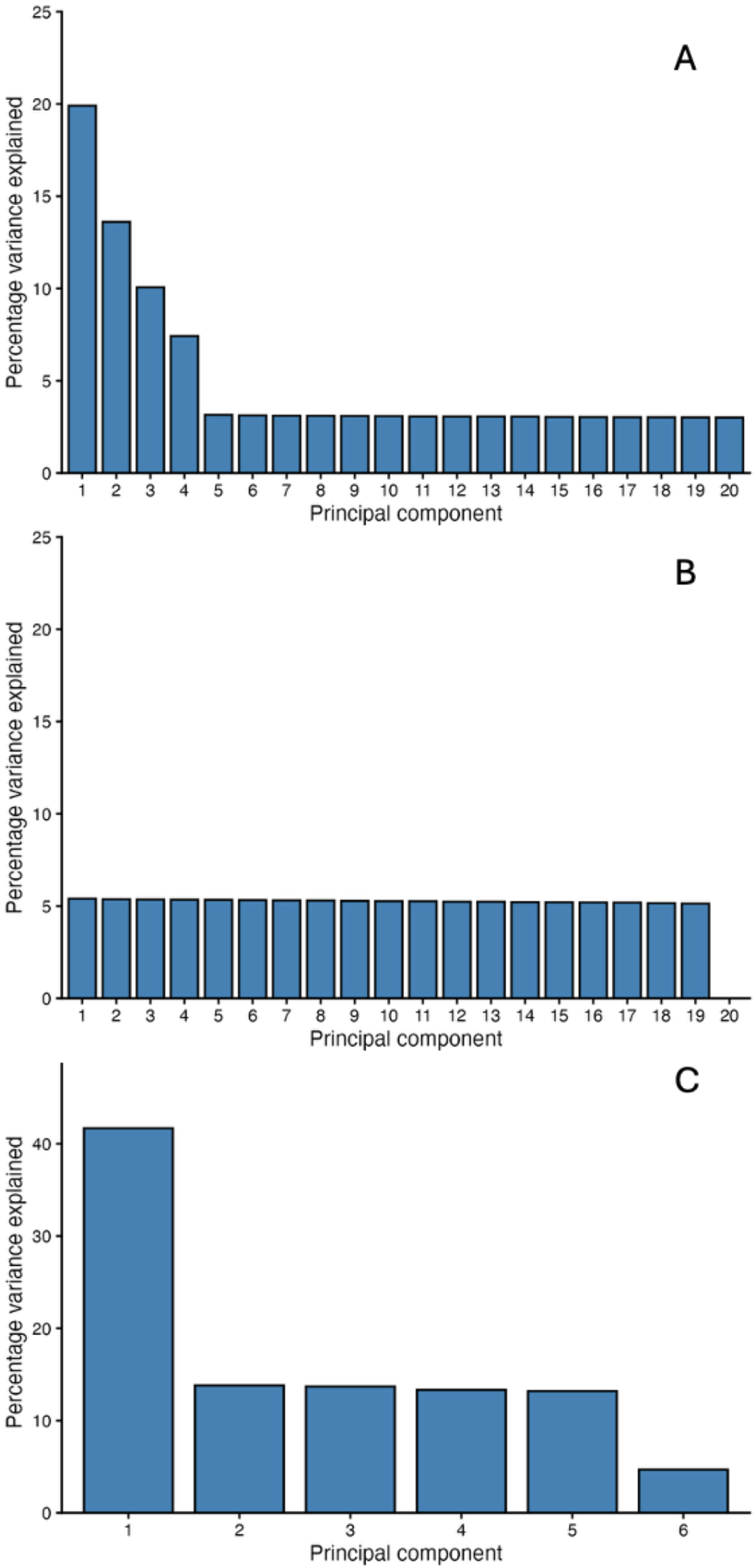
Percent variance explained (PVE) of each PC from PCA. (A) PVE of PCA including all samples. (B) PVE of PCA including only Isabela samples. (C) PVE of PCA including only eastern island samples. Related to Figure 2.

**Figure S7.**
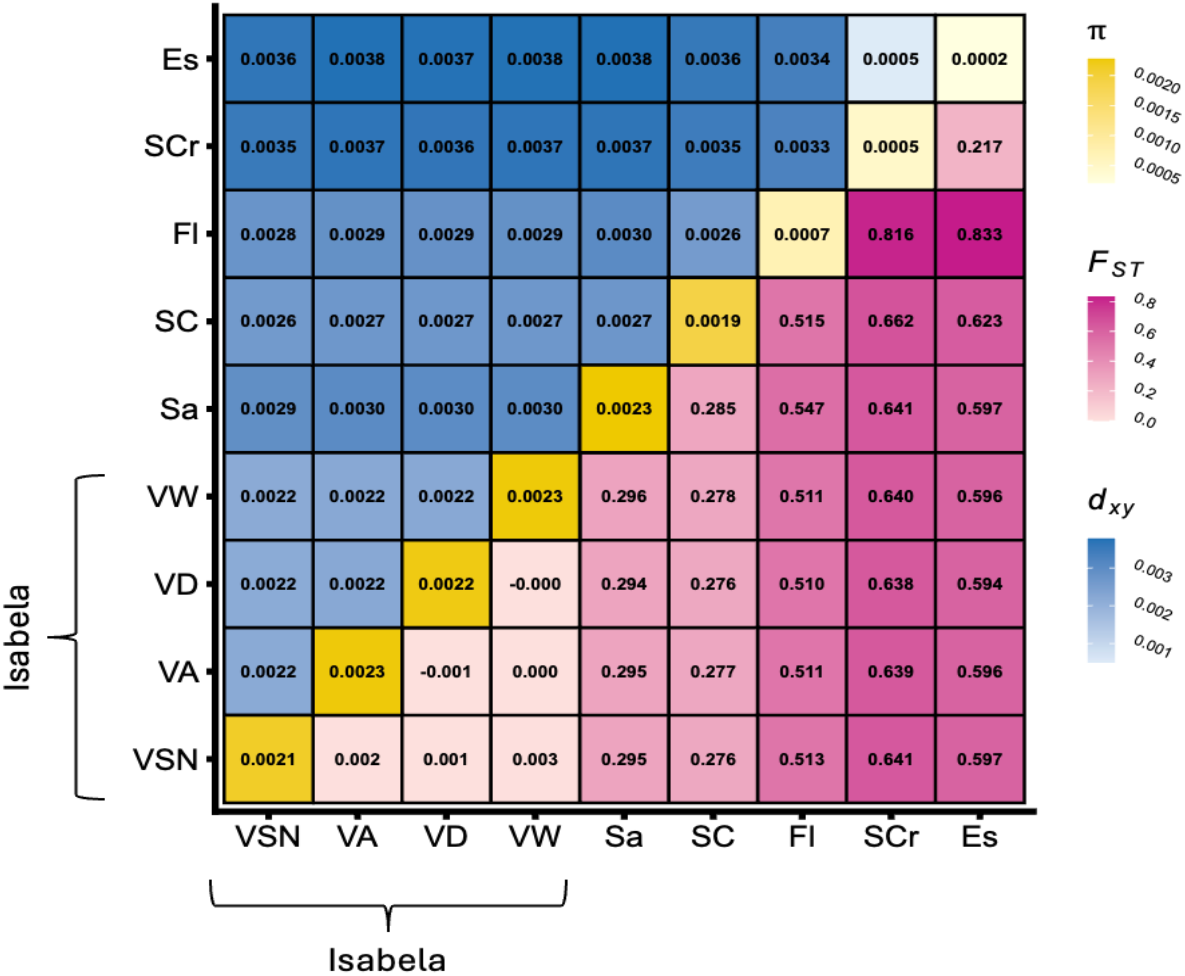
Genetic structure matrix including pairwise comparisons with each volcano of Isabela separately. Related to Figure 2.

**Figure S8.**
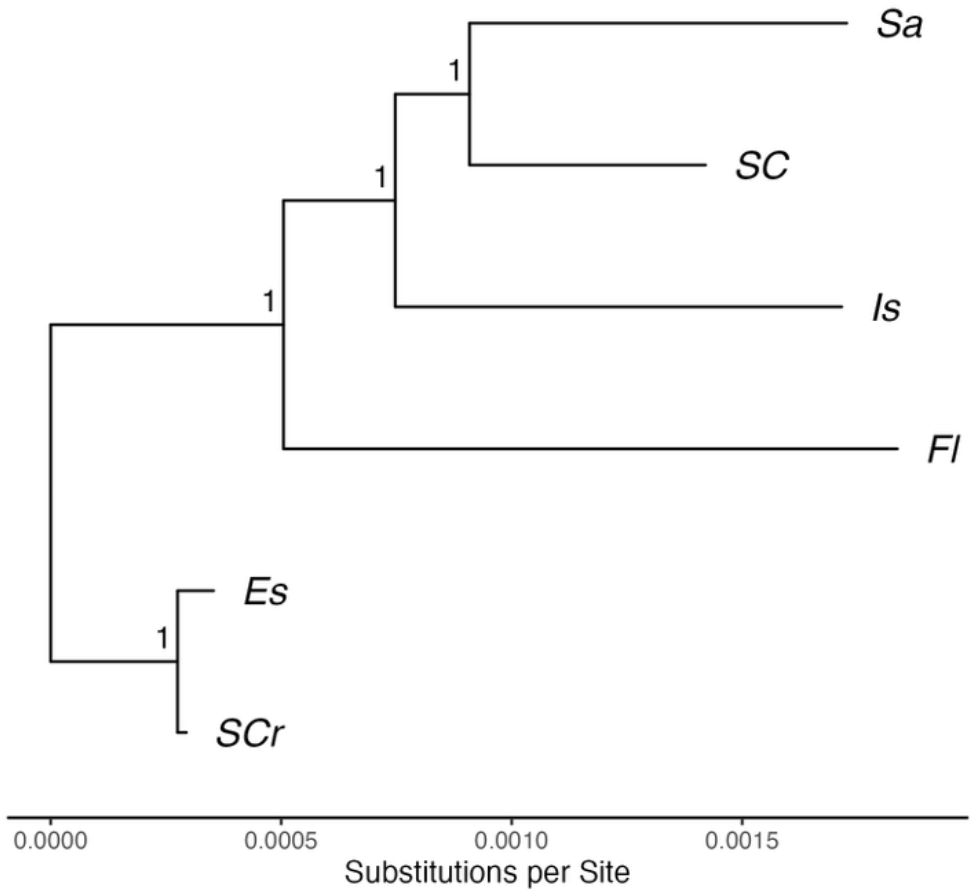
ASTRAL-IV species tree topology estimated from 8,848 gene trees with branch lengths estimated with CASTLES-II. Related to Figure 4.

**Figure S10:**
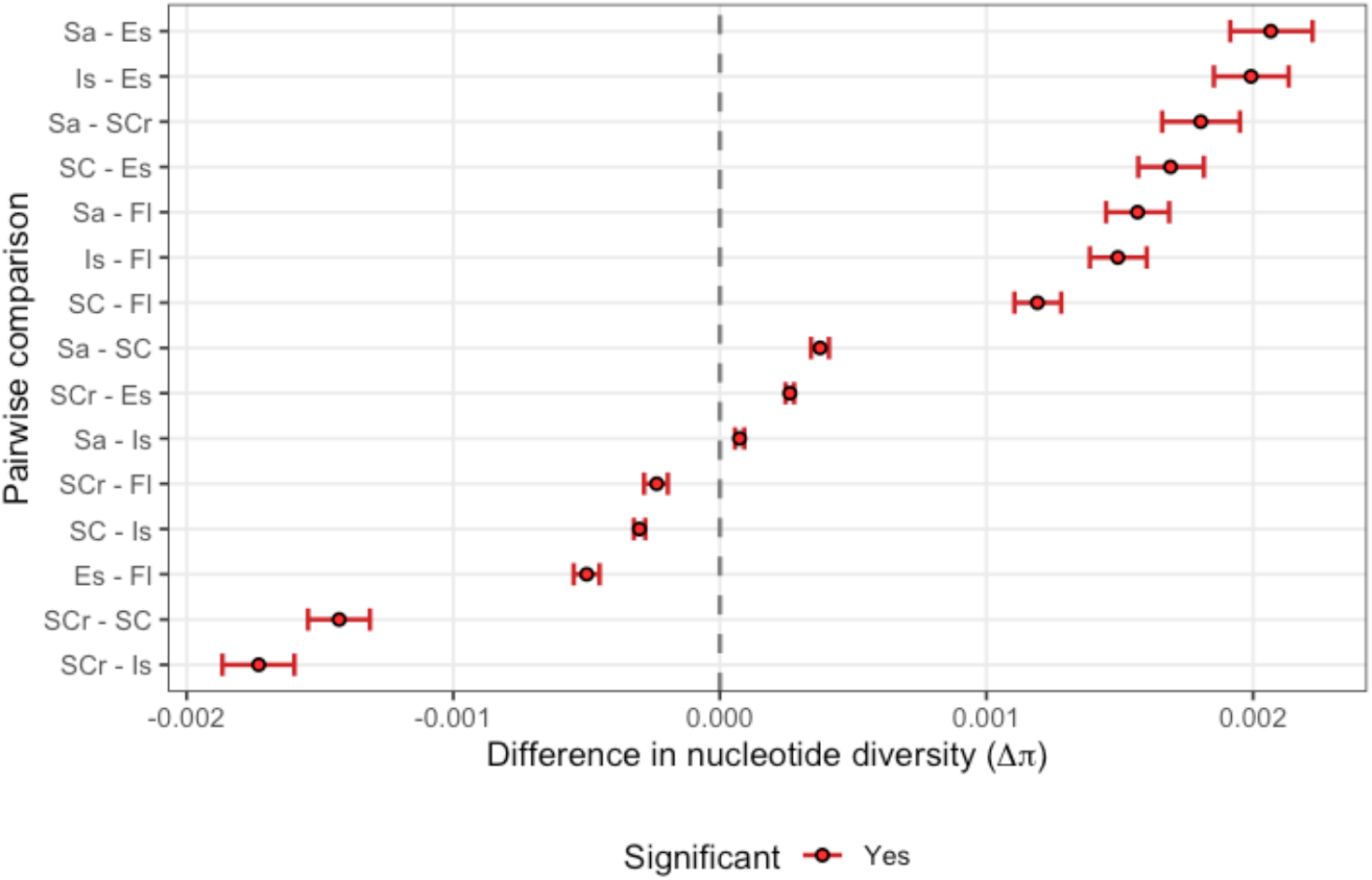
Differences in genome-wide nucleotide diversity calculated for all pairwise comparisons. Points represent mean bootstrap estimates of the pairwise difference in nucleotide diversity and error bars represent 95% confidence intervals from 1,000 replicates of a 1 Mbp genomic block bootstrap. Comparisons were considered significantly different when the CI did not overlap zero.

**Figure S11.**
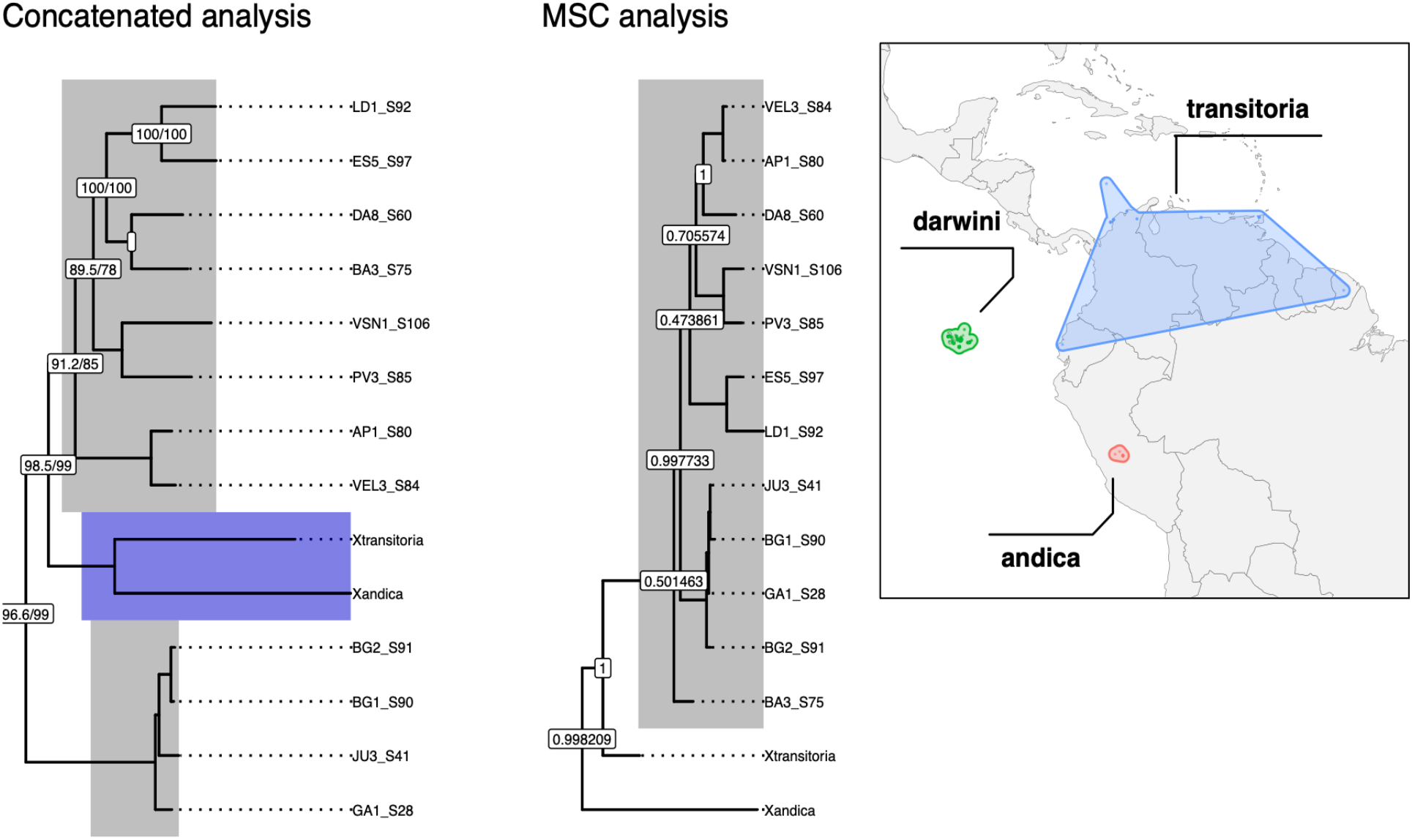
Relationships among Galápagos taxa and mainland taxa based on 440 UCE loci. (Left) Maximum likelihood tree based on a concatenated alignment. Node labels represent bootstrap support values. (Middle) Species tree estimated with ASTRAL-IV using 440 UCE gene trees. (Right) Map showing the extant distribution of the mainland South American sister taxa.

**Table S1.**
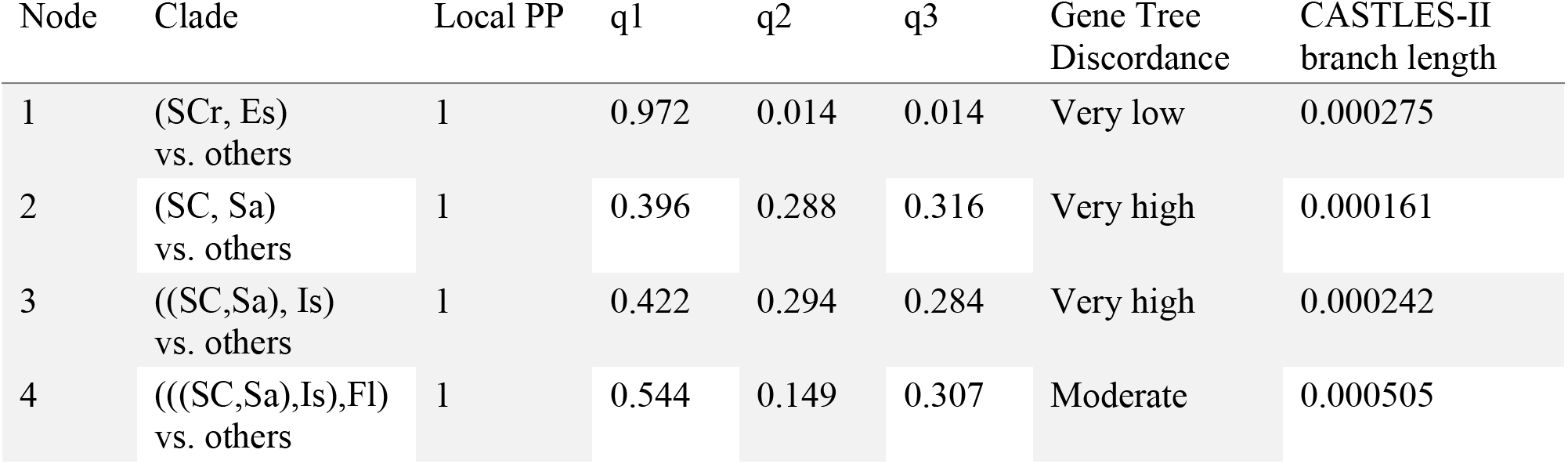
Quartet frequencies (q1–q3) and support values for internal nodes of the species tree. Local posterior probabilities (local PP) provide node support. q1 denotes the proportion of gene trees supporting the inferred topology, while q2 and q3 represent alternative topologies. Lower q1 values indicate substantial gene tree discordance, consistent with incomplete lineage sorting or rapid diversification. Related to Figure 3.

**Table S1.**
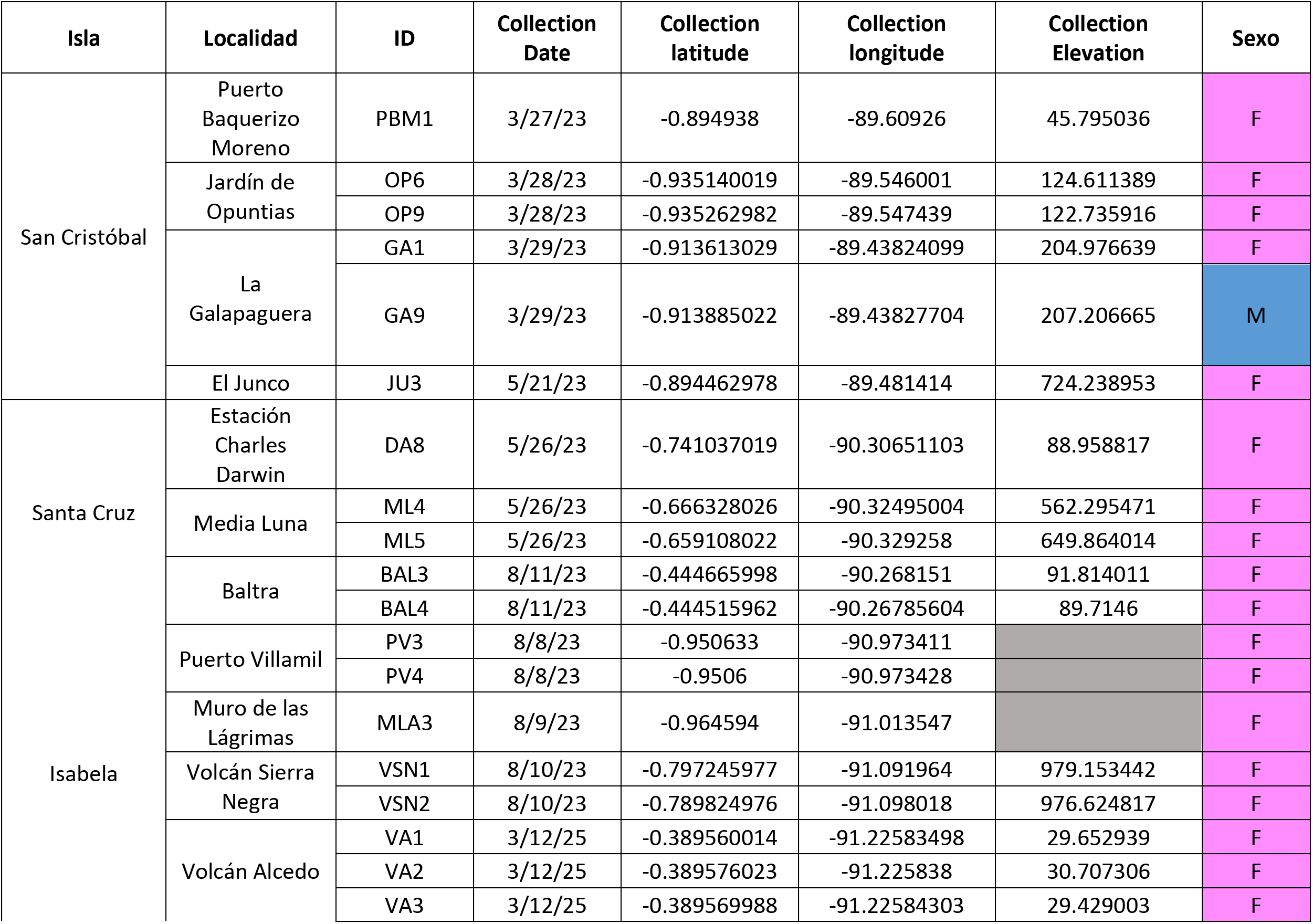
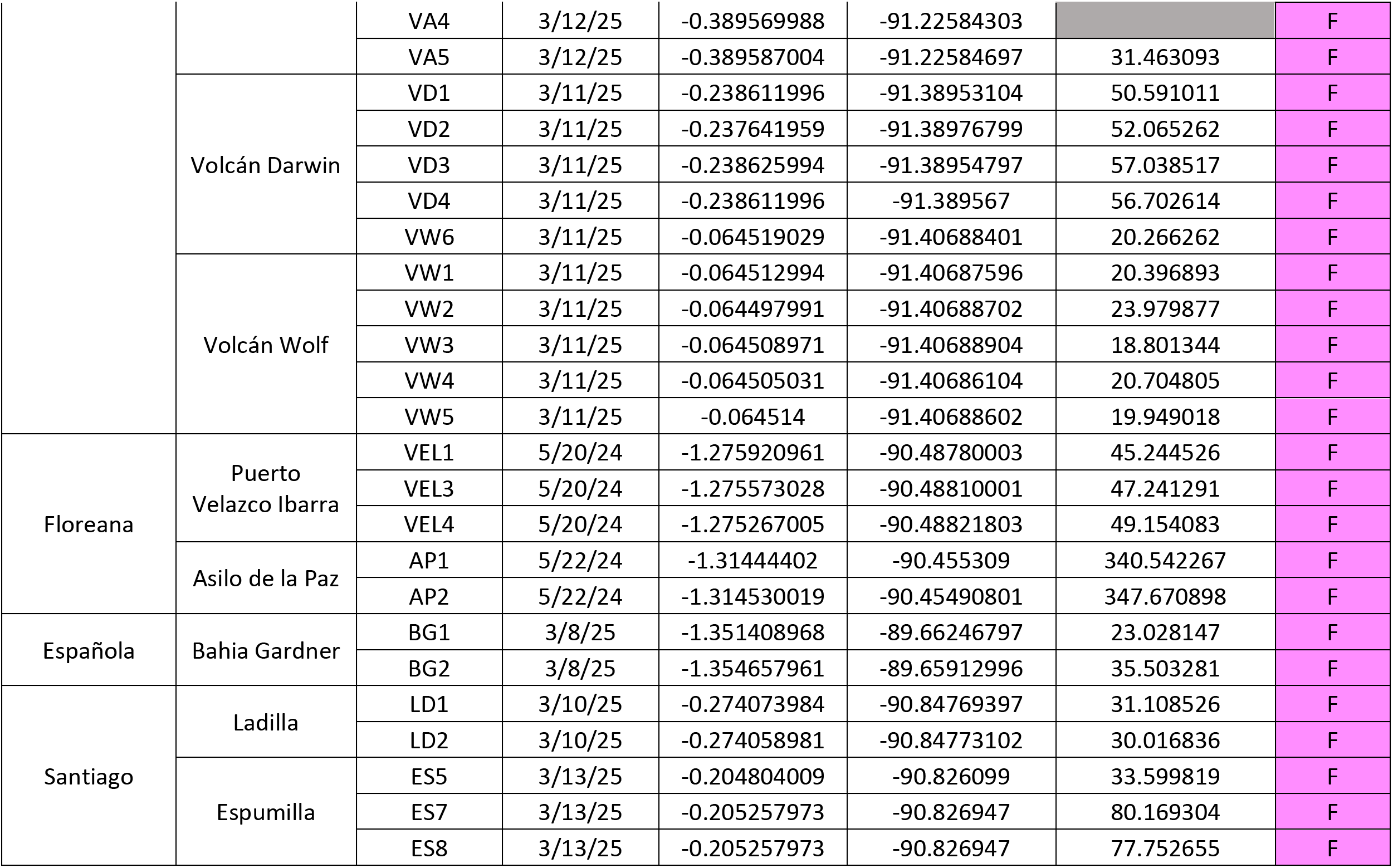
Sampling details of the bee specimens used in this study.

## Notes

### Competing Interest Statement

The authors have declared no competing interest.

### Summary of Updates

Wording in abstract and discussion modified to improve clarity; minor citation corrections in introduction; acknowledgments updated; supplemental files updated

